# Contrasting wood chemical traits are associated with differences in wood degradation and esca susceptibility among grapevine cultivars

**DOI:** 10.64898/2025.12.09.693160

**Authors:** Pierre gastou, Thomas carayol, Gwenaëlle comont, Nathalie ferrer, Nathalie boizot, Claudia rouveyrol, Pierre pétriacq, Chloé E. L. delmas

## Abstract

Deciphering the interplay between microbial communities and host defence mechanisms is key to understanding plant health. In perennial plants, the balance between endophytes and wood (*i.e.* secondary xylem) defence responses governs wood degradation and vascular disease expression. Esca is a complex vascular disease contributing to grapevine decline, with an incidence variable across cultivars but the mechanisms underlying its expression and varietal susceptibility remain unclear. We assessed relationships between internal wood degradation and esca foliar symptoms in *Vitis vinifera* L. cultivars grown in a common garden, and, at the cultivar level, we tested for correlations between (i) susceptibility to wood decay and esca expression (n = 16 cultivars) and (ii) wood biochemical traits, and healthy wood endophytic microbial communities (n = 23 cultivars). Unlike other types of necrosis, white-rot necrotic wood was significantly more abundant in plants that had expressed esca leaf symptoms in previous years, particularly in the most susceptible cultivars. These cultivars also contained significantly lower levels of constitutive wood extractives. However, glycosylated phenylpropanoids accumulated in the wood of esca-symptomatic plants, especially in highly susceptible cultivars. By contrast, esca expression and varietal susceptibility had only a marginal effect on the diversity, composition and putative functions of microbial communities in healthy wood. They did not influence either the relative abundance of *Fomitiporia mediterranea*, the putative causal agent of white-rot in grapevine. Esca susceptibility appears primarily linked to wood degradability and metabolic responses, rather than healthy wood’s microbial communities, suggesting that the use of less susceptible varieties together with white-rot removal might attenuate grapevine decline.

## Introduction

The wood (*i.e.* secondary xylem) of perennial plants is involved in long-term physiological function and molecular dialogue with xylem-dwelling microorganisms (Yadeta and Thomma, 2013; Torres-Ruiz et al., 2024). Its biochemical properties and the dynamics of wood-endophytic microorganism interactions over time, across organs and plant genotypes, may therefore directly (through plant defence responses or abundances of pathogens or their antagonists) or indirectly (through defence priming or physiological modulations) affect the expression of complex vascular diseases. These diseases are major causes of woody plant decline worldwide and are characterised by physical (*e.g.* wood necrosis) and physiological (*e.g.* hydraulic dysfunctions, impaired carbon storage and growth) damage, and symptom expression, sometimes distal to infected areas (Agrios, 2005; Yadeta and Thomma, 2013). Changes in the endophytic microbiome have been reported but are less studied than the edaphic and aerial compartments (Darriaut et al., 2021; Gómez-Aparicio et al., 2022; Del Frari et al., 2025). They include increases in pathogen abundance and activity, particularly putative virulence factor production (Broberg et al., 2018; Nerva et al., 2022; Chambard et al., 2026), reduced microbial diversity (Steinrucken et al., 2016), and losses of beneficial microbial functions (Arnault et al., 2023). These modifications are not documented in all pathosystems (Raghavendra et al., 2017; Del Frari et al., 2019). Conversely, specific physicochemical defence mechanisms operate in wood. Anatomical barriers limit the axial, lateral and radial spread of pathogens (Morris et al., 2016) and are reinforced by active responses, such as vascular occlusion, the impregnation of wood structures with lignin and the accumulation of secondary metabolites (Morris et al., 2016, 2020; Silva et al., 2020), potentially primed by specific microbial taxa (Liu et al., 2017).

Significant and broad intraspecific variability has been reported for susceptibility to vascular diseases involved in woody plant decline (Rosado et al., 2010; Venturas et al., 2013). The genetic diversity of the microbiome and plant responses to pathogens underlies varietal susceptibility to these diseases. Plant genotype directly influences the diversity, composition and ecological function of wood-inhabiting communities through nutritional, physical (e.g. wood composition) and chemical (e.g. plant-pathogen molecular interactions) selection pressures, therefore modulating disease progression (Lamit et al., 2014; Gomes et al., 2019; Mina et al., 2020). Genotypes also differ in their physiological and molecular responses to microbial colonisation: occlusion formation, lignin deposition (Niza et al., 2015; Silva et al., 2020), induction of metabolic pathways involved in defence (Khattab et al., 2021; Gomes et al., 2023; Patanita et al., 2025). However, the role of wood properties and physiology in woody-plant interactions with vascular pathogens remains largely unknown, despite these traits probably underlying genotypic variability in susceptibility to dieback.

Esca disease of grapevine is a relevant model, as this complex syndrome contributes to decline in grapevine (*Vitis vinifera* L.), a woody crop of high socio-economic value. Esca is characterised by the summer expression of interveinal leaf scorch symptoms (chlorotic and necrotic phenotypes) known as *tiger-stripe*s (Gramaje et al., 2018). They may be accompanied by bunch wilting or sudden canopy desiccation (*i.e.* apoplectic phenotypes), ultimately leading to plant death (Li et al., 2017; Gastou et al., 2024). Leaf symptom onset is associated with the appearance of a brown stripe of non-conducting tissue along the xylem (Lecomte et al., 2012, 2024; Gastou et al., 2026), and systemic physiological disturbances (Bortolami et al., 2019, 2021a,b; Dell Acqua et al., 2024, 2025; Chambard et al., 2026). Symptomatic vines typically display decreases in healthy wood area in trunks and cordons, with increases in internal necrosis enriched in wood pathogens — white-rot necrosis (mostly enriched in Basidiomycota) and black necrosis (mostly enriched in Ascomycota) (Maher et al., 2012; Bruez et al., 2020; Gastou et al., 2026). However, such necroses also occur in asymptomatic vines (*i.e.* that have never expressed esca foliar symptom; Gastou et al., 2026) and their role in symptom expression across grapevine varieties remains unclear. The direct role of wood pathogens in symptom expression is increasingly called into question (Hofstetter et al., 2012; Del Frari et al., 2019; Gastou et al., 2026; Monod et al., 2025).

Explorations of the intraspecific diversity of wood microorganisms may shed light on the mechanisms underlying esca expression and varietal susceptibility in grapevine. The susceptibility of *Vitis vinifera* L. to esca (*i.e.* the varietal mean incidence estimated by the proportion of plants exhibiting leaf symptoms and declining phenotypes across years) varies considerably between cultivars (Etienne et al., 2024; Gastou et al., 2024). The putative mechanisms underlying these differences include a greater accumulation of specific defence metabolites and lower water use efficiency in highly susceptible genotypes (Gastou et al., 2024; Gastou et al., 2025). The role of internal wood degradation in shaping varietal susceptibility remains insufficiently explored, although studies on narrow diversity panels suggest a possible role for wood composition (Rolshausen et al., 2008), metabolic defences (Lemaitre-Guillier et al., 2020), microbial community dynamics (Bekris et al., 2021), and pathogen aggressiveness (Laveau et al., 2009).

We investigated the factors affecting wood-endophytic microorganism interactions in a range of mature grapevine cultivars in a common garden. We studied the relationships between internal wood integrity and esca leaf symptom expression, and the plant and microbial traits underlying grapevine susceptibility to wood degradation and esca expression. We hypothesised that (i) the extent of symptomatic internal necrosis would be greater in plants with a history of esca symptoms, particularly for the most susceptible cultivars; (ii) less susceptible cultivars are constitutively richer in lignin, defence metabolites and micro-organisms identified as antagonists of wood pathogens, whereas highly susceptible cultivars exhibit greater wood pathogen abundance; (iii) the plant response to esca involves similar metabolic mechanisms in all cultivars but is more intense in highly susceptible genotypes.

## Materials and methods

### Origin of the plant material

We sampled material from 23 *V. vinifera* L. cultivars (Supplementary Table S1) from the VitAdapt common garden experimental vineyard, at the *Institut National de Recherche pour l’Agriculture, l’Alimentation et l’Environnement* (INRAE) research station (Villenave d’Ornon, Nouvelle-Aquitaine, France), at 44°47’23.83 N’’, 0°34’39.3’ W’. The plants were grown in a randomised block design with homogeneous management, as described by Gastou et al. (2024). Vines were trained in a double Guyot system, with a trunk and two permanent arms from which annual stems are pruned. These arms carry pruning wounds, often associated with dry wood areas known as pruning-wound necrosis.

All vines were individually monitored for the incidence of esca foliar symptoms and declining phenotypes each summer from 2017 to 2023 (Gastou et al., 2024), and again in 2024. Sampling was performed within three blocks with comparable incidences of esca foliar symptoms (blocks 1, 3 and 5; Gastou et al., 2024). We selected cultivars along the varietal gradient of esca foliar symptom incidence. We defined *varietal esca susceptibility* as the mean annual proportion of symptomatic plants per cultivar over the 2017–2024 period, averaged across experimental blocks.

### Whole-vine sampling, cross-sectioning and wood sampling for phenotyping wood integrity and biochemical composition

We sampled 96 individual plants from 16 cultivars during dormancy between 5th November and 22nd November 2024 (Supplementary Fig. S1A). We cut 55 control plants (*i.e.* with an asymptomatic history since 2017; between two and six plants per cultivar; 16 cultivars) and 41 plants with a history of esca symptoms (*i.e.* at least one year with esca leaf symptoms between 2017 and 2024; one to four plants per cultivar; 14 cultivars) 25 cm above the ground with a chainsaw. On the same day, four 2 cm-thick transversal cross-sections were cut with a mitre saw: two from the trunk (8 cm and 15 cm from the vine head) and one from each arm (8 cm from the vine head). Sections were photographed with a D34000 camera (Nikon, Tokyo, Japan) for the visual quantification of wood types. Apparently healthy wood samples (3 g of wood per cross-section) were collected from upper trunk sections with a wood chisel for analyses of wood chemical composition. Apparently healthy wood is visually identified as white-yellow wood with a solid structure and no visible signs of decay (necrosis, punctuation). All equipment and the sapwood surface were disinfected with 96% ethanol [v/v] between samplings. Samples were placed directly in liquid nitrogen and ultimately stored at -80°C.

### Visual quantification of wood types

Cross-sections (*n* = 386) were analysed with ImageJ 1.53t (Wayne Rasband and contributors, National Institutes of Health; https://imagej.net/software/fiji/), using a protocol adapted from Maher et al. (2012). We first measured the total area and circumference of each cross-section. We then separately measured the areas of apparently healthy wood, black necrotic wood, white-rot necrotic wood and dry wood (Fig.1A and B). Yellowish-brown areas that could not be classified as an identifiable type of necrosis were considered to be transition zones. We quantified the visually altered sapwood perimeter by measuring the portion of the sapwood (outer xylem ring; McElrone et al., 2021) that is visually identified as white-rot necrotic wood, black necrotic wood, transition zone or dry wood. We attributed a qualitative index (from 0 to 3) for black punctate necrosis to each cross-section based on two-by-two comparisons of the density of black punctuations (Supplementary Fig. S1B).

**Fig. 1.**
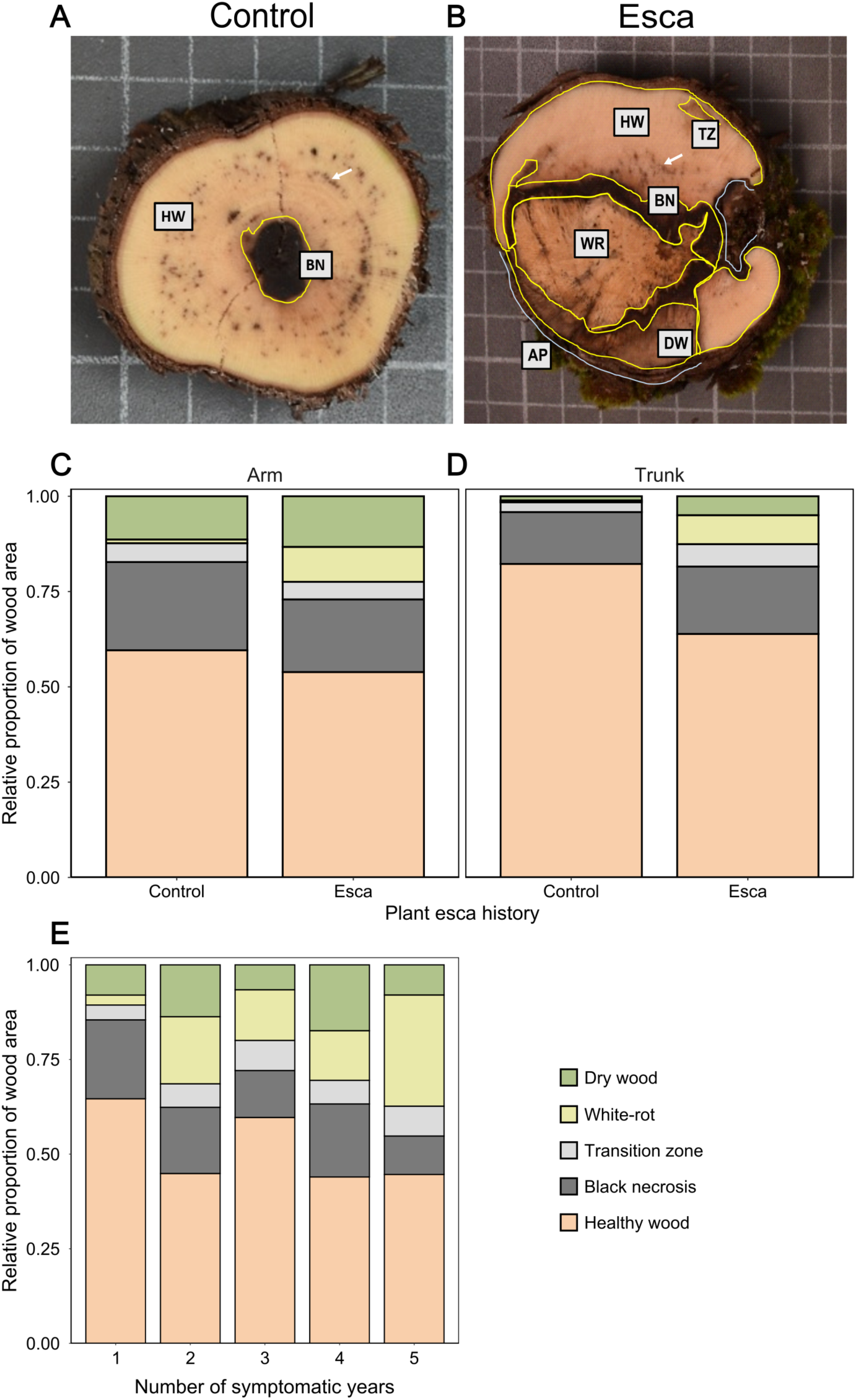
Distribution of healthy and necrotic wood in the trunk, for different organs and plant esca histories. **A, B,** Grapevine trunk cross-sections showing the different wood types quantified. Healthy wood (HW), black necrotic wood (BN), white-rot necrotic wood (WR), transition zones (TZ) and dry wood (DW) areas are circled in yellow. The visually altered sapwood perimeter (AP) is shown in light blue. The white arrow indicates necrotic wood with black punctuations. The control plant (Petit Verdot) in (A) displays only a narrow central area of black necrotic wood and a large area of healthy wood containing black punctuations. The plant that previously displayed esca symptoms (Cabernet-Sauvignon) in (B) has diverse large necrotic areas, altering the sapwood perimeter. **C, D, E,** Mean relative proportions of healthy wood, black necrotic wood, transition zones, white-rot and dry wood according to plant esca history in (C) arm (*n* = 191 sections) and (D) trunk samples (*n* = 195 sections), and (E) according to the number of years of symptoms in plants previously displaying esca symptoms (*n* = 164 sections). The bars are coloured by wood type. Detailed statistics are provided in Supplementary Table S3.

### Determination of wood chemical composition

Wood cell-wall composition was determined by analytical biochemistry (*n* = 96 samples from 16 cultivars). Analyses were performed on 300 mg of ground and sieved wood powder (75-500 µm). The proportion of extractive components was determined by sequential extractions with 96% boiling ethanol to eliminate phenolic acids, non-structural carbohydrates, waxes, pigments, then with boiling water to eliminate tannins and proteins (Sluiter et al., 2005; TAPPI T 204 om-88 standard, 1987). The wood powder free from extractives was hydrolysed with 5 ml of 72% sulfuric acid for 2h at room temperature under agitation. A second hydrolysis was performed with 3% sulfuric acid during 1 hour at 120°C. After filtration through a glass funnel filter with porosity 3, the hydrolysis gave insoluble residue used for Klason lignin dosage and soluble filtrate used for both soluble lignin and cellulose and hemicellulose determination. Soluble lignin was quantified by spectrophotometry at 205 nm (TAPPI standard UM250, 1991) whilst insoluble lignin (Klason lignin) was quantified by weighing the residue after washes with water to achieve a neutral pH and a dry at 105 °C for 16 h (TAPPI standard T 222 om-02, 2002). Total lignin content was expressed as the sum of insoluble Klason and soluble lignin. The proportions of cellulose and hemicellulose were determined indirectly by using high-performance liquid chromatography (HPLC) to quantify the monomeric sugars released by hydrolysis: glucose from cellulose, and xylose, arabinose, mannose and galactose from hemicellulose (Gao et al., 2014). To do this, 25 ml of filtered hydrolysate was neutralized with 1 g of calcium carbonate during 16h at 4°C then clarified with Chromafil Xtra PET-20/25 0.2µm (Macherey-Nagel). 1 mL was dried under vacuum and resuspended in 100 μL ultra-pure water containing 2 mg mL−1 mannitol as an internal standard. 25 µL were injected into a Chromaster high-performance liquid chromatography system (VWR Hitachi) equipped with a Rezex™ RPM-Monosaccharide Pb+2 (8%) column (Phenomenex), then eluted with ultrapure water at a flow rate of 0.6 ml/min. Monomeric sugars were quantitatively detected with an evaporative light scattering detector (ELSD) 85 (VWR Hitachi with the following parameters: Temperature 60°C; Gain 5; Offset 0; Filter 6 s; Sampling time 100 ms-10 Hz; Nitrogen pressure 3.5 bar), and the peak areas were electronically integrated using OpenLAB CDS EZChrom (Agilent). Quantities of monomeric sugars were calculated from the calibration curves obtained with 10, 20, 40 and 60 µg of standard sugars (Sigma).

### Field sampling of healthy wood for metabolomics and metabarcoding

In addition to the whole-vine sampling described above, we collected 123 trunk samples from 23 cultivars on different individual plants from the same vineyard, between 24th July and 7th August 2023. Wood was collected from 78 asymptomatic plants (*i.e.* no esca foliar symptoms at sampling; at least 3 plants par cultivar) between two and six plants per cultivar and 47 symptomatic plants (*i.e.* tiger-stripe esca foliar symptoms at sampling; 3 plants per cultivar were used where possible).

Sapwood samples were collected with a PRO 18V Cordless percussion drill (AEG, Frankfurt am Main, Germany) from the upper section of the trunk, at a depth of 2 cm after bark removal (Supplementary Fig. S1A). The pieces of wood were collected in a Petri dish. Only apparently healthy wood was retained, placed in a 5 ml microtube, which was immediately immersed in liquid nitrogen. All equipment and the sapwood surface were disinfected with 96% ethanol [v/v] before the collection of each sample. Samples were ultimately stored at -80°C for metabarcoding and metabolomic analyses.

### Untargeted metabolomics

Healthy (non-necrotic) wood samples (*n* = 123) were ground in liquid nitrogen with a one-ball mill TissueLyser II (Qiagen, Hilden, Germany). We then sent 10mg of lyophilised wood tissue to the MetaboHUB-Bordeaux Metabolome facility (https://metabolome.u-bordeaux.fr/; Villenave-d’Ornon, France) for untargeted metabolomics by the LCMS profiling of semi-polar extracts. Metabolites were extracted with robots, as previously described (Luna et al., 2020; Dussarrat et al., 2022), in a solvent containing 80% ethanol [v/v], 0.1% formic acid [v/v] and supplemented with methyl vanillate as an internal standard (250 µg/mL). Each sample was extracted twice by sonication (15 min, ice-cold water), the supernatants were pooled, and extracts were filtered through 0.22 µm MultiScreen GV filtration plates (Merck, Molsheim, France). Quality controls (prepared by pooling 50 µl from each sample) and extraction blanks were included on each plate.

Untargeted metabolic profiling was performed by UHPLC-LTQ-Orbitrap mass spectrometry (LCMS) with an Ultimate 3000 ultra-high-pressure liquid chromatography (UHPLC) system coupled to an LTQ-Orbitrap Elite mass spectrometer interfaced with an electrospray ionisation source (ESI, Thermo Fisher Scientific, Bremen, Germany) operating in negative-ion mode. Chromatographic separation was achieved on a Gemini C18 reverse-phase column (150 × 2 mm, 3 µm; Phenomenex) at 40°C using a water/acetonitrile gradient containing 0.1% formic acid. Mass spectra were acquired at 30,000 resolution in data-dependent acquisition mode, and QC samples were injected every ten samples to monitor analytical stability. Raw LCMS data were processed in MS-DIAL v.4.9 with optimised parameters, yielding 4,758 features, which were then curated (signal-to-noise ratio > 10; coefficient of variation in quality controls < 30%), yielding 3,035 features. Annotations were performed as described by Dell’Acqua et al. (2025) with both FragHUB (Dablanc et al., 2024) and in-house standard databases. Annotated features were compliant with MSI confidence levels ranging from MSI2 to MSI4 (Sumner et al., 2007). The extraction protocol targets a broad range of semi-polar compounds, including phenolics and other secondary metabolites. Given that metabolomic analyses were performed on independent samples and that only a subset of detected features could be annotated (MSI levels 2–4), the relative contribution of specific metabolite classes to the total extractive fraction (quantified as described below) could not be quantified.

### Fungal and bacterial DNA metabarcoding

#### DNA extraction

We extracted genomic DNA from ground wood tissues (60 mg, *n* = 123) with the DNeasy Plant Mini Kit (Qiagen, Hilden, Germany) according to the manufacturer’s instructions. We included 17 extraction blanks (*i.e.* reagents without sample). Extracted double-stranded DNA was quantified with a DS-11 spectrophotometer (DeNovix, Wilmington, USA) and the concentration of all samples was standardised to a maximum of 15 ng/µl.

#### Targeted DNA amplification

Fungal DNA was amplified by targeting the ITS region of the fungal rRNA operon with the ITS1f/ITS2 primer pair (CTTGGTCATTTAGAGGAAGTAA/ GCTGCGTTCTTCATCGATGC). Bacterial DNA was amplified by targeting the V5/V6 region of the 16S RNA gene with the chloroplast-excluding primer pair 799f/1115r (AACMGGATTAGATACCCKG/AGGGTTGCGCTCGTTG). Primers were provided by Eurogentec, Seraing, Belgium. Amplifications were performed independently, with the protocol described below.

For each sample, polymerase chain reaction (PCR) mixture consisted of template genomic DNA (2 µl), 2X Platinum Hot Start PCR Master Mix (12 µl; Invitrogen, Waltham, USA), 2.5µl of each primer (forward and reverse), and nuclease-free water (5.5µl; Invitrogen, Waltham, USA). Amplifications were performed as follows: initialisation at 95°C for 2 min; 35 cycles of denaturation at 95°C for 45 s, annealing at 60°C for 1 min, extension at 72°C for 1 min 30; and a final extension at 72°C for 10 min. The quality and specificity of the amplification were assessed by electrophoresis in 2% agarose (m/v) TAE gels. We included 30 PCR blanks (*i.e.* without DNA template) for each amplification and 56 positive controls (11 pure DNA extracts each of *Yamadazyma oceani* and *Yamadazyma barbieri* for ITS amplification, presented in Fournier et al., 2025; 8 pure extracts of *Bacillus thuringiensis* and 26 pure extracts of *Erwinia persicina* for 16S amplification).

#### DNA sequencing

We sent PCR products to the Genome Transcriptome Facility of Bordeaux (Cestas, France) for Illumina MiSeq sequencing (v3 chemistry, 2×300 bp). PCR product purification, multiplex identifiers and sequencing adapter addition, library sequencing and sequence demultiplexing (with exact index search) were performed independently for ITS and 16S amplicons by the sequencing service.

#### Bioinformatics analyses

All bioinformatic analyses were performed with the FROGS pipeline on the Galaxy server (https://vm-galaxy-prod.toulouse.inrae.fr/galaxy_main/; Escudié et al., 2018). Data underwent preprocessing (*i.e.* merging, filtering and dereplicating) and single-linkage SWARM clustering was performed on sequences with an aggregation distance of 1. PCR chimeras were removed from each sample, and OTUs were filtered with the minimum prevalence method and a minimum proportion of 5e-05.

OTUs underwent BLAST taxonomic annotation with ITS UNITE Fungi 8.3 and 16S SILVA 138.1 pintail 100, for ITS and 16S data, respectively. The affiliation of each OTU was checked manually. If the BLAST affiliation was ambiguous or inconsistent with RDP affiliation, we obtained BLAST affiliations from the NCBI (nr/nt) nucleotide collection (https://blast.ncbi.nlm.nih.gov/Blast.cgi?PROGRAM=blastn&PAGE_TYPE=BlastSearch&LINK_LOC=blasthome) and, for ITS data, trunkdiseaseID.org (https://www.grapeipm.org/d.live/?q=td-lab-dna). We tried to maximise consensus but NCBI was considered to be the reference database if disagreement persisted.

### Data analysis

#### Statistical tools

All analyses were performed in *R* v.4.2.1. Modelling procedures were performed with the *lme4* package. Pearson’s tests were performed and visualised with the *corrplot* package. Metabarcoding data were studied with the *phyloseq* and *vegan* packages. Metabolomic data were analysed in *Metaboanalyst* v.6.0 and *R* v.4.2.1. All tests were performed with a significance level of 5% (α = 0.05). All models were checked graphically for the normality of residuals (QQ-plot) and homogeneity of the residual variance (residuals *vs*. fitted, residuals *vs*. predictors). All pairwise post-hoc comparisons were performed with Tukey’s tests.

#### Modelling wood integrity and composition

We first investigated the effects of organ, cultivar, plant esca history and their interactions on the overall distribution of wood types in a permutational analysis of variance (PERMANOVA) with Euclidean distances. We then investigated the effects of organ, cultivar, plant esca history and their interactions on the log1p-transformed relative proportions of each wood type (*i.e.* healthy wood, black necrotic wood, white-rot necrotic wood, transition zones, dry wood) and of the visually altered sapwood perimeter, with independent linear mixed models (LMM). We investigated the effects of cultivar, organ, plant esca history, and their interactions on the ordinal index of black punctate necrosis with a cumulative link mixed model (CLMM). In plants with a prior history of esca symptoms, we investigated the effects of plant health status (2024) on wood type proportions. We also investigated the effect of the number of symptomatic years.

We assessed Pearson’s correlations between (i) the relative proportion of each wood type, the proportion of visually altered sapwood perimeter, the index of black punctate necrosis and (ii) varietal esca susceptibility for each cultivar, a quantitative variable defined as the mean incidence of foliar symptoms per cultivar over the 2017–2024 period. We performed the same analyses for the distribution of wood components: PERMANOVA for their overall distribution, independent LMM procedures for each wood component, and Pearson’s correlations with varietal esca susceptibility.

Finally, we investigated the effects of the relative proportion of each wood component on the log1p-transformed proportions of healthy wood, white-rot necrotic wood, black punctate necrosis, transition zones, dry wood and visually altered sapwood perimeter with independent LMM procedures. The ordinal index of black punctate necrosis was modelled with a CLMM. We compared effect sizes with computed standardised estimates of the model parameters. We also analysed Pearson’s correlations between (i) the relative proportion of each wood component and (ii) the mean proportion of white-rot necrotic wood in esca plants.

#### Analyses of metabolomics and metabarcoding data

##### Varietal susceptibility clusters

For multivariate factorial analyses of omics datasets (metabarcoding and metabolomics), we defined cultivar susceptibility groups based on both foliar symptom incidence and white-rot necrotic wood proportion. These two variables capture complementary aspects of esca expression at the cultivar level, allowing an exploratory approach to identify putative wood-microorganism interactions underlying white-rot and esca susceptibility. K-means clustering was performed with mean esca foliar symptom incidence per cultivar (2017-2024) and mean white-rot necrotic wood proportion per cultivar in plants with a history of esca foliar symptoms (2024, computed in this study), with 10 random initialisations. The number of clusters was evaluated using silhouette analysis, which indicated that both k = 2 and k = 3 provided meaningful structures. k = 3 was retained to distinguish contrasting extreme susceptibility profiles (less and highly susceptible cultivars) and intermediate ones (moderately susceptible cultivars). Finally, we obtained the following varietal esca susceptibility classes: low (*n* = 10 cultivars), moderate (*n* = 5 cultivars) and high (*n* = 8 cultivars) (Supplementary Fig. S2).

##### Trunk healthy wood metabolome

Normalisation against the sample median, square root data transformation and Pareto scaling were applied to metabolomic data before each analysis. We first assessed the effects of cultivar, varietal esca susceptibility class (based on both foliar symptoms 2017-2024 and white-rot necrotic wood proportion 2024), and plant health status on trunk metabolome structure by PERMANOVA.

We then specifically evaluated the effect of plant health status on trunk metabolome structure by PERMANOVA. We identified differentially abundant (fold change > 2 and a *p*-value < 0.01, no False Discovery Rate correction) metabolites for each pairwise comparison by Volcano plot analyses. Putative contaminants (*i.e.* non-plant metabolites) were manually curated based on published findings and our own knowledge. Chemical taxa were assigned to differentially abundant chemical features based on InChiKeys, with the structural ontology tool ClassyFire (Djoumbou Feunang et al., 2016) and the ClassyFire Batch interface (https://cfb.fiehnlab.ucdavis.edu/). We performed graphical analyses to compare enrichment in specific structural ontologies for each pairwise comparison.

A similar process was performed independently to evaluate the effect of varietal esca susceptibility class in plants with a history of esca symptoms. A similar process was performed independently to evaluate the effect of varietal esca susceptibility in asymptomatic plants (i.e. constitutive metabolome).

We also evaluated differences in metabolomic variance between plant health statuses by performing a test for homogeneity of multivariate dispersions (PERMDISP), with Euclidean distances and 999 permutations. Pairwise post-hoc comparisons on mean distances to centroids were performed for significant global tests.

##### Microbial communities in healthy trunk wood

All analyses were performed separately for fungal and bacterial data. Before analysing metabarcoding data, we decontaminated the dataset with the package metabaR (Zinger et al., 2021). Contaminant OTUs were identified based on their relative abundance in negative controls and removed from the dataset. Additional filtering steps included the removal of non-target taxa and OTUs with low sequence similarity to reference databases (<80%). Samples with low sequencing depth (<1,000 reads) or high contamination levels (>10% contaminant reads) were also not further analysed. Tag-jump artefacts were mitigated by applying a relative abundance threshold of 0.01 at the sample level, which reduced spurious diversity in negative controls while preserving biological signal in environmental samples.

We first assessed the effects of cultivar, varietal esca susceptibility class and plant health status on the diversity and structure of healthy trunk microbial (*i.e.* fungal or bacterial) communities. We compared Shannon diversity index between susceptibility classes and plant health statuses in LMM procedures. Effects on community structure were then assessed by PERMANOVA on CLR-transformed data increased by a pseudocount of 1. The multivariate homogeneity of dispersions was checked before the performance of pairwise comparisons.

A similar process (*i.e.* Shannon index, PERMANOVA) was performed independently to evaluate the effect of varietal esca susceptibility in plants with esca symptoms. We classified microbial OTUs into three functional groups: putative grapevine wood pathogens (fungi only), putative antagonists of grapevine wood pathogens (fungi and bacteria), and other taxa (fungi and bacteria). This taxonomic classification to genus level was based on the available scientific literature (as detailed in Gastou et al., 2026). We investigated the effects of varietal esca susceptibility on the relative proportions of these functional classes by ANOVA. Moreover, we performed a differential analysis with the ALDEx2 procedure on RLE-transformed data with a pvalue_cutoff = 0.05, to identify the microbial species most likely to explain differences between varietal esca susceptibility classes.

A similar process (*i.e.* Shannon index, PERMANOVA, functional classification, ALDEx2) was performed independently to evaluate the effect of varietal esca susceptibility in asymptomatic plants (*i.e.* constitutive microbial communities).

## Results

### Effects of cultivar and plant esca history on wood integrity

Cultivar, organ (trunk vs. arm of the vine), and plant esca history significantly influenced the overall distribution of wood types (Supplementary Table S2). Wood type differences between plant esca histories (control *vs*. esca) depend on the organ (Fig. 1C and D) and cultivar (Fig. 2; Supplementary Table S2). The differences between arms and trunks are detailed in Supplementary Table S2 and mostly consisted of a larger proportion of dry wood in arm samples (Fig. 1C and D).

**Fig. 2.**
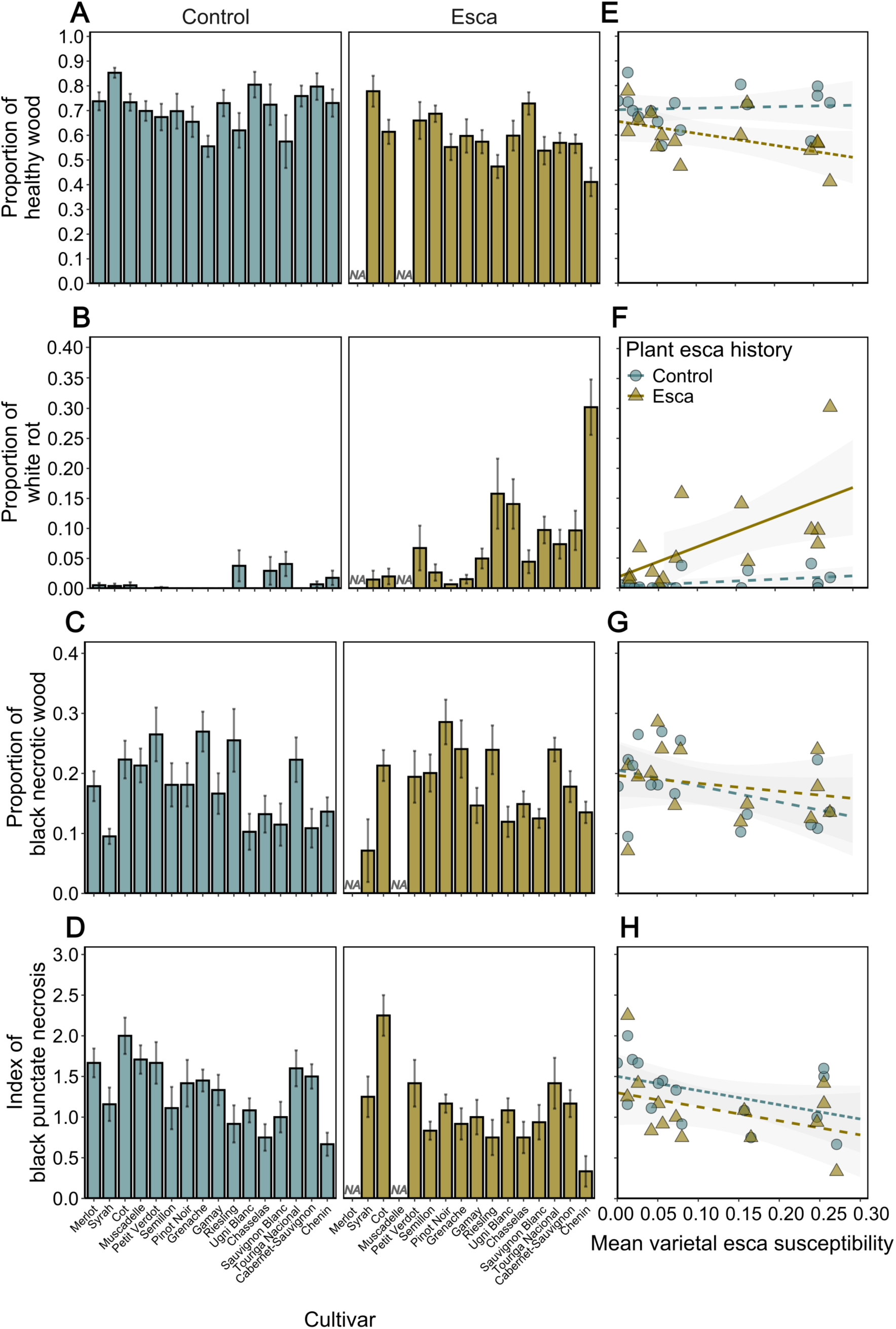
Effect of cultivar on wood-type proportions and relationship to varietal mean esca susceptibility, by variety. **A, B, C, D,** Mean proportions of (A) healthy wood, (B) white-rot necrotic wood, (C) black necrotic wood, and (D) index of black punctate necrosis (mean ± SEM) across cultivars for control (*n* = 222 sections, *n* = 55 plants, *n* = 16 cultivars) plants and plants with a history of esca foliar symptoms (*n* = 164 sections, *n* = 41 plants, *n* = 14 cultivars). *P*-values correspond to cultivar effects in linear mixed models fitted separately for control and esca plants (A, B), or on the entire dataset (C, D). **E, F, G, H,** Varietal relationships between esca susceptibility (foliar symptoms 2017-2024) and mean proportions of (E) healthy wood, (F) white-rot necrotic wood, (G) black necrotic wood, and (H) index of black punctate necrosis. Each dot corresponds to the value measured for a specific cultivar. Correlation coefficients and associated *p*-values for Pearson’s tests are provided in the main text. Detailed statistics are provided in Supplementary Table S3. Blue dots/bars correspond to control plants, brown triangles/bars correspond to plants with a history of esca foliar symptoms. Significant relationships are indicated by solid lines, marginally significant relationships by dashed lines, and non-significant relationships by dotted lines. Wood-type distribution is averaged across two trunk sections and two arm sections per plant.

Plant esca history had a significant effect on the proportion of healthy wood (*P* = 10^-5^) with plants with a history of esca foliar symptoms having a mean of 17% less healthy wood than control plants (Table 1). In plants with a history of esca foliar symptoms, the proportion of white-rot necrotic wood was 12 times higher and the proportion of visually altered sapwood perimeter was almost twice as high (Supplementary Fig. S4A) than in control plants (Table 1). The proportions of dry wood and black necrotic wood and the index of black punctate necrosis were not significantly affected by plant esca history (Table 1; Supplementary Fig. S4B).

**Table 1.** Comparisons of wood-type distribution between previously esca-symptomatic plants and controls. The mean and standard error are shown for each wood type. Comparisons were made through independent linear mixed modelling procedures. ns: non-significant. *p* < 0.1, * *p* < 0.05, ** *p* < 0.01, *** *p* < 0.001.

| Wood type | Control<br>($n = 55$ plants) | Previously esca-symptomatic plants<br>($n = 41$ plants) | Comparison |
| --- | --- | --- | --- |
| Healthy wood | $0.71 \pm 0.01$ | $0.59 \pm 0.02$ | *** |
| Black necrotic wood | $0.18 \pm 0.009$ | $0.18 \pm 0.009$ | ns |
| Transition zones | $0.04 \pm 0.003$ | $0.05 \pm 0.005$ | . |
| White-rot necrotic wood | $0.007 \pm 0.002$ | $0.08 \pm 0.01$ | *** |
| Dry wood | $0.06 \pm 0.009$ | $0.09 \pm 0.01$ | ns |
| Visually altered sapwood perimeter | $0.11 \pm 0.01$ | $0.21 \pm 0.02$ | *** |
| Index of black punctate necrosis | $1.38 \pm 0.06$ | $1.04 \pm 0.06$ | ns |

In plants with a history of esca foliar symptoms, the number of symptomatic years (ranging from 1 to 5; Fig. 1E) significantly affected the proportions of most wood types, except black necrotic wood and dry wood (Supplementary Table S3). The proportion of healthy wood was significantly higher for plants with one year of symptoms than for those with two, but did not differ significantly from that in plants with three or more symptomatic years (Fig. 1E). Conversely, the proportions of white-rot necrotic wood and visually altered sapwood perimeter were significantly higher for plants with two years of symptoms than in those with one, but did not differ significantly from that in plants with three or more years of symptoms (Fig. 1E). Wood-type distribution was similar in plants with one year of symptoms and in control plants (Fig. 1E). Interestingly, 29.3% (*n* = 12 of 41) of plants with a history of esca symptoms from nine cultivars had no white-rot necrotic wood. All these plants with a symptomatic history but no white-rot had a history of only one year of esca symptoms, either in the summer preceding the sampling (three plants) or before (nine plants). The maximal mean proportion of white-rot necrotic wood in plants with esca symptoms was 0.45 (Chenin with a history of symptoms in five years). Conversely, none of the control plants had mean white-rot necrotic wood proportions exceeding the mean value in plants with esca (0.08).

Cultivar significantly affected the proportions of healthy wood (Fig. 2A; Supplementary Table S3), black necrotic wood (Fig. 2C) and transition zones, the index of black punctate necrosis (Fig. 2D), but not the proportions of dry wood and visually altered sapwood perimeter. Cultivar significantly affected the proportion of white-rot necrotic wood in plants with a history of esca foliar symptoms (*P =* 0.02) but not in control plants (*P =* 0.11; Fig. 2B). The mean proportion of white-rot necrotic wood per cultivar in plants with a history of esca foliar symptoms ranged from less than 0.01 (Pinot Noir) to 0.30 ± 0.05 (Chenin).

At cultivar level, varietal mean esca susceptibility tended to decrease with increasing mean proportion of healthy wood in plants with a history of esca foliar symptoms (*r =* -0.50, *P =* 0.07) but not in control plants (Fig. 2E; Supplementary Table S3). It was positively correlated with the proportion of white-rot necrotic wood in plants with a history of esca foliar symptoms (*r =* 0.62, *P =* 0.02) but not in control plants (*r =* 0.22, *P =* 0.41; Fig. 2F). It tended to increase with increasing proportion of visually altered sapwood perimeter in plants with a history of esca foliar symptoms but not in control plants, and to decrease with increasing proportion of black necrotic wood (Fig. 2G) and index of black punctate necrosis (Fig. 2H). It was not correlated with the proportions of dry wood and transition zones.

Finally, in plants with a history of esca foliar symptoms, plant health status in 2024 (*i.e.* asymptomatic *vs.* symptomatic in the summer preceding the sampling) did not significantly affect the distribution of wood types (Supplementary Table S4).

### Variations of wood composition across cultivars with different susceptibilities to esca

The mean overall proportion of wood components across *V. vinifera* L. cultivars was: 14 ± 0.2% extractives, 15 ± 0.1% Klason lignin, 3 ± 0.04% soluble lignin, 47 ± 0.3% cellulose and 20 ± 0.1% hemicellulose.

The overall distribution of wood components was significantly influenced by cultivar, but not plant esca history or their interaction (Fig. 3A and B; Supplementary Table S5). More specifically, cultivar significantly influenced the relative proportions of all wood components studied (Fig. 3A and B).

**Fig. 3.**
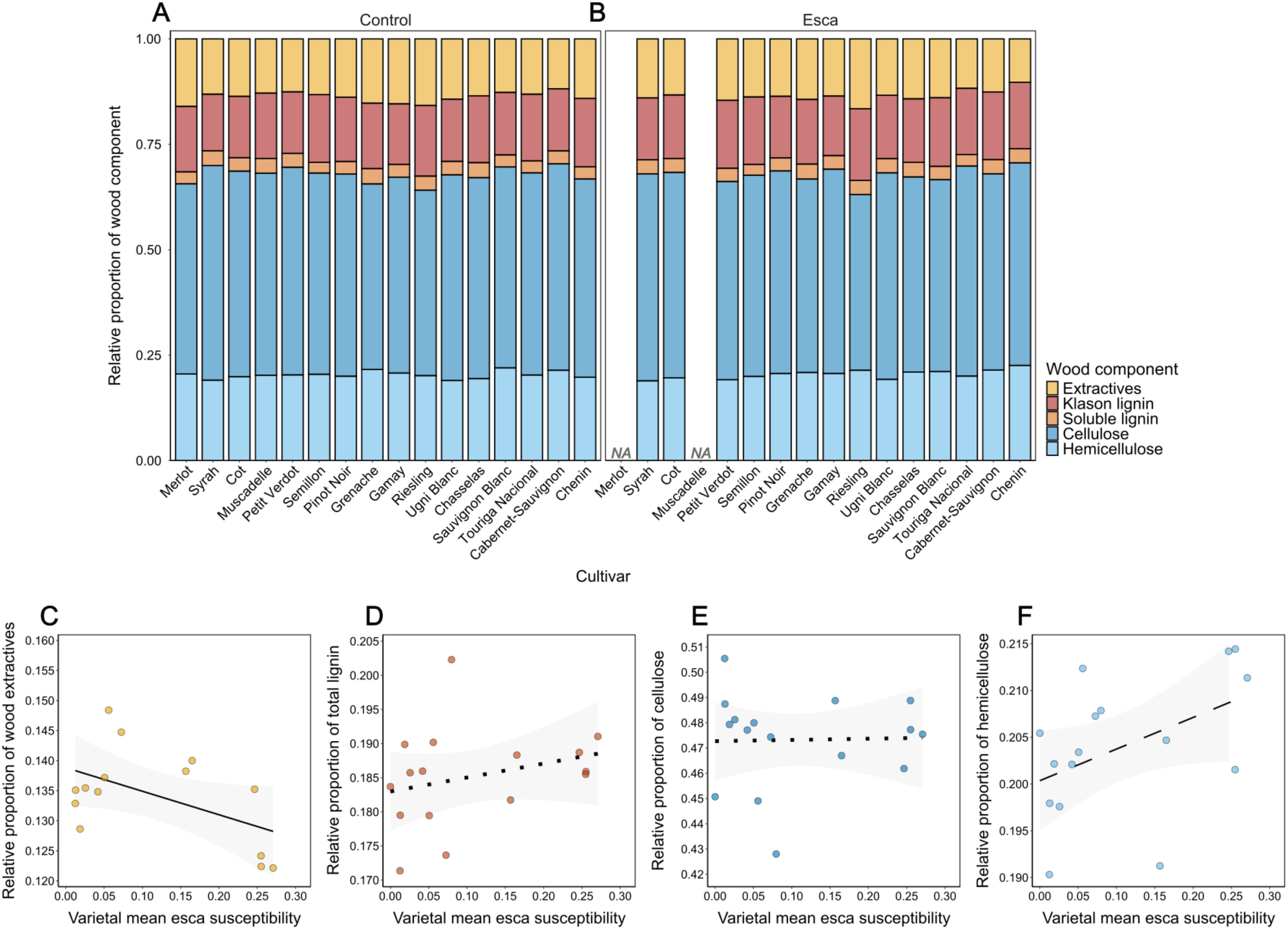
Distribution of wood components in upper trunk sections across plant esca histories and cultivars and relationship with varietal esca susceptibility. **A, B,** Mean relative proportions of extractives, Klason lignin, soluble lignin, cellulose and hemicellulose across cultivars in (A) control plants (*n* = 55 plants, *n* = 16 cultivars) and (B) plants with a history of esca foliar symptoms (*n* = 41 plants, *n* = 14 cultivars). **C, D, E, F,** Varietal relationships between varietal esca susceptibility (foliar symptoms 2017-2024) and mean proportions of (C) extractives, (D) total lignin (*i.e.* Klason and soluble lignins), (E) cellulose and (F) hemicellulose. Each dot corresponds to the value measured for a specific cultivar. Correlation coefficients and associated p-values in Pearson’s tests: r = -0.51, *P* = 0.04 for extractives; r = 0.28, *P* = 0.29 for total lignin; r = 0.03, *P* = 0.92 for cellulose; r = 0.46, *P* = 0.07 for hemicellulose. Detailed statistics are provided in Supplementary Table S6. Significant relationships are shown by solid lines, marginally significant relationships by dashed lines, and non-significant relationships by dotted lines. Bars and dots are coloured by wood component.

At cultivar level, varietal mean esca susceptibility was significantly negatively correlated with the mean proportion of wood extractives (*r =* -0.51, *P =* 0.04; Fig. 3C). It was not significantly correlated with the mean proportion of Klason lignin (*r =* 0.32, *P =* 0.22), soluble lignin (*r =* -0.13, *P =* 0.63), total lignin (*r =* 0.28, *P =* 0.29; Fig. 3D) or cellulose (*r =* 0.03, *P =* 0.92; Fig. 3E). It tended to increase with the mean proportion of hemicellulose (*r =* 0.46, *P =* 0.07; Fig. 3F).

### Relationships between wood composition and white-rot necrotic wood proportion

The effect of wood composition on the proportion of white-rot necrotic wood was assessed in upper trunk sections, with cultivar, plant esca history and their interaction as covariates (Supplementary Table S6).

At plant level, the proportion of white-rot necrotic wood decreased significantly with increasing proportion of extractives in plants with a history of esca foliar symptoms (β = -0.09, *P =* 0.04; Fig. 4A) while it was not significantly influenced by the proportions of total lignin (Fig. 4B), or cellulose (Fig. 4C).

**Fig. 4.**
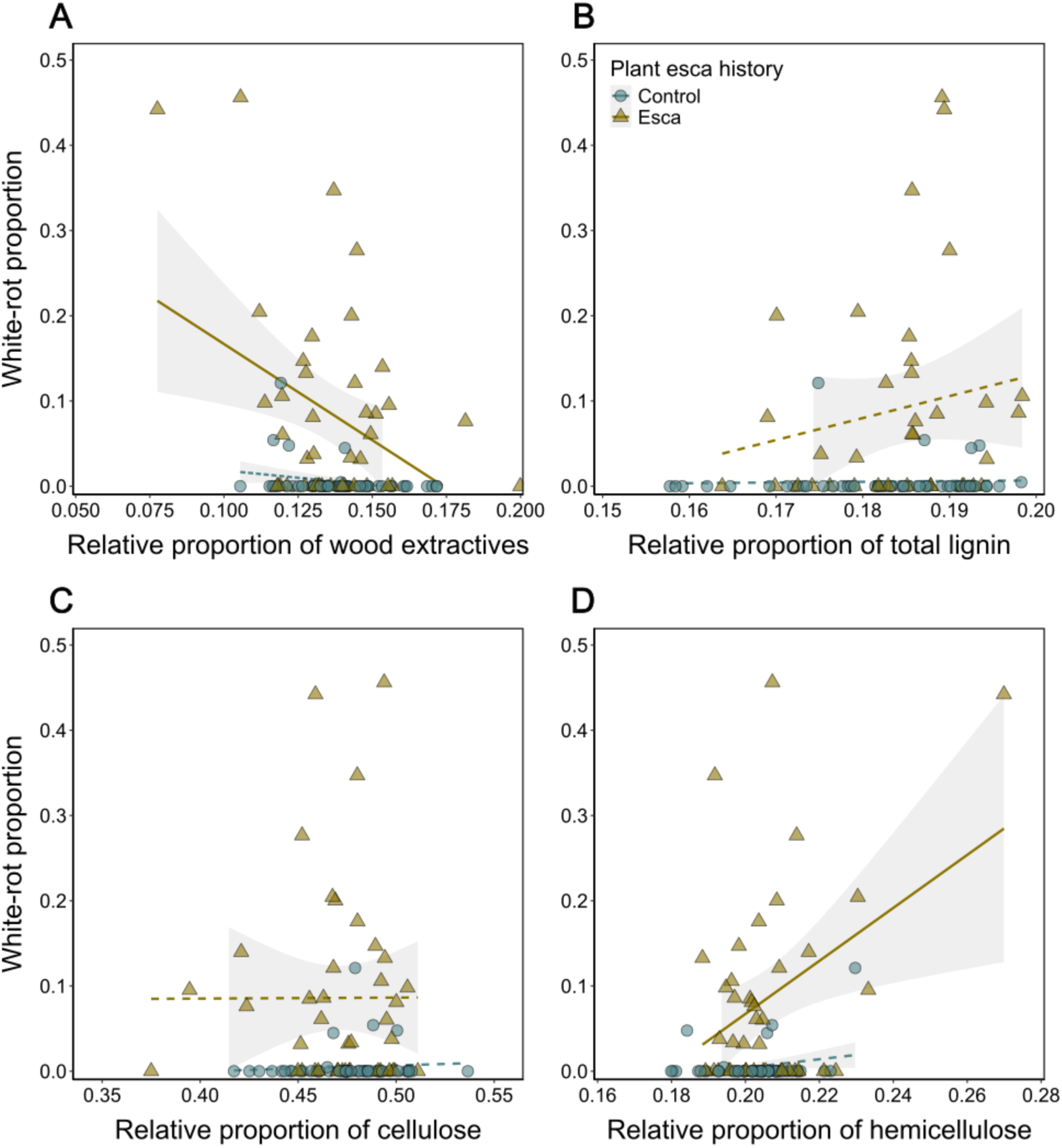
Relationship between wood composition and white-rot necrotic wood proportion in upper trunk sections of control plants (*n* = 55 plants, *n* = 16 cultivars) and plants with a history of esca foliar symptoms (*n* = 41 plants, *n* = 14 cultivars). **A,** Mean proportion of extractives. **B,** Mean proportion of total lignin (*i.e.* Klason and soluble lignins). **C,** Mean proportion of cellulose. **D,** Mean proportion of hemicellulose. Each dot corresponds to the value for an individual plant. Blue dots/bars correspond to control plants, brown triangles/bars correspond to esca plants. Significant relationships are indicated by solid lines, marginal relationships by dashed lines, and non-significant relationships by dotted lines. Detailed statistics are presented in Supplementary Table S7.

Interestingly, the proportion of white-rot necrotic wood increased significantly with increasing hemicellulose proportion at the plant level (β = 0.04, *P =* 0.04; Fig. 4D), but this relationship was driven by a single plant (previously esca-symptomatic Chenin) with an extreme phenotype: very high proportions of white-rot and hemicellulose.

At cultivar level, mean white-rot necrotic wood proportion per cultivar in plants with a history of esca foliar symptoms was positively correlated with the proportion of hemicellulose (*r =* 0.74, *P =* 0.002), and marginally negatively correlated with the proportion of extractives (*r =* 0.49, *P =* 0.07). It was not significantly correlated with the proportions of Klason lignin (*r =* 0.34, *P =* 0.34), soluble lignin (*r =* 0.21, *P =* 0.47), total lignin (*r =* 0.41, *P =* 0.14) or cellulose (*r =* -0.18, *P =* 0.54).

### Effects of cultivar and esca expression on the trunk metabolome

We quantified 3,035 metabolic features across 123 trunk samples. To identify the factors influencing metabolic variations, we first compared the trunk metabolome across plant esca health statuses and varietal esca susceptibility classes (based here on mean foliar symptom incidence and white-rot necrotic wood proportion). Plant health status and varietal susceptibility, but not their interaction (Supplementary Table S7) influenced overall trunk metabolic structure slightly but significantly. Multivariate dispersion differed significantly between asymptomatic and symptomatic plants (*P* = 0.004; Supplementary Fig. S5A). Mean distance to the centroid was higher for symptomatic plants (30.93) than asymptomatic plants (28.18). Plants with esca symptoms displayed enrichment in a larger number of compounds than asymptomatic plants (30 *vs.* 13 features; Supplementary Fig. S5B; see *Asymptomatic vs Symptomatic ALL comparison* in Supplementary Table S7). An enrichment in “phenylpropanoids and polyketides” (flavonoids, isoflavonoids, and depsides or depsidones) was observed in symptomatic plants (6 *vs.* 1 feature; Supplementary Fig. S5C; Supplementary Table S7).

To understand how esca susceptibility level modulates induced wood metabolism, we analysed plants with esca symptoms separately. We found that varietal esca susceptibility significantly influenced trunk metabolic structure (Fig. 5A). No clear difference was noted in the number of compounds displaying enrichment between less and moderately susceptible cultivars (15 *vs.* 18 features; Fig. 5B; *see Low vs Moderate S comparison* in Supplementary Table S7). Enrichment was observed for a larger number of compounds in highly than in less (32 *vs.* 13 features; Fig. 5C; *see Low vs High S comparison* in Supplementary Table S7) and moderately (24 *vs.* 6 features; Fig. 5D; 5C; *see Moderate vs High S comparison* in Supplementary Table S7) susceptible cultivars. A clear enrichment in “phenylpropanoids and polyketides” was observed in moderately susceptible cultivars relative to less susceptible cultivars (6 *vs.* 2 features; Fig. 5E), and this enrichment was even stronger in highly susceptible than in less susceptible cultivars (10 *vs.* 2 features; Fig. 5F; Fig. 6). Glycosylated compounds (*e.g.* flavonoid-glycosides as quercitrin, phenolic glycosides as 2-(beta-D-Glucopyranosyloxy)benzyl 3-(hexopyranosyloxy)-6-hydroxy-2-methoxybenzoate, stilbene glycosides as 4-[(E)-2-(3-Hydroxy-5-methoxyphenyl)vinyl]phenyl beta-D-glucopyranoside; putative annotations) were overrepresented among the phenylpropanoids for which enrichment was detected in moderately and highly susceptible cultivars (Fig. 6). Enrichment in “lipids and lipid-like compounds” was found in highly susceptible cultivars relative to moderately susceptible cultivars (5 *vs.* 1 features; Fig. 5G; Fig. 6).

**Fig. 5.**
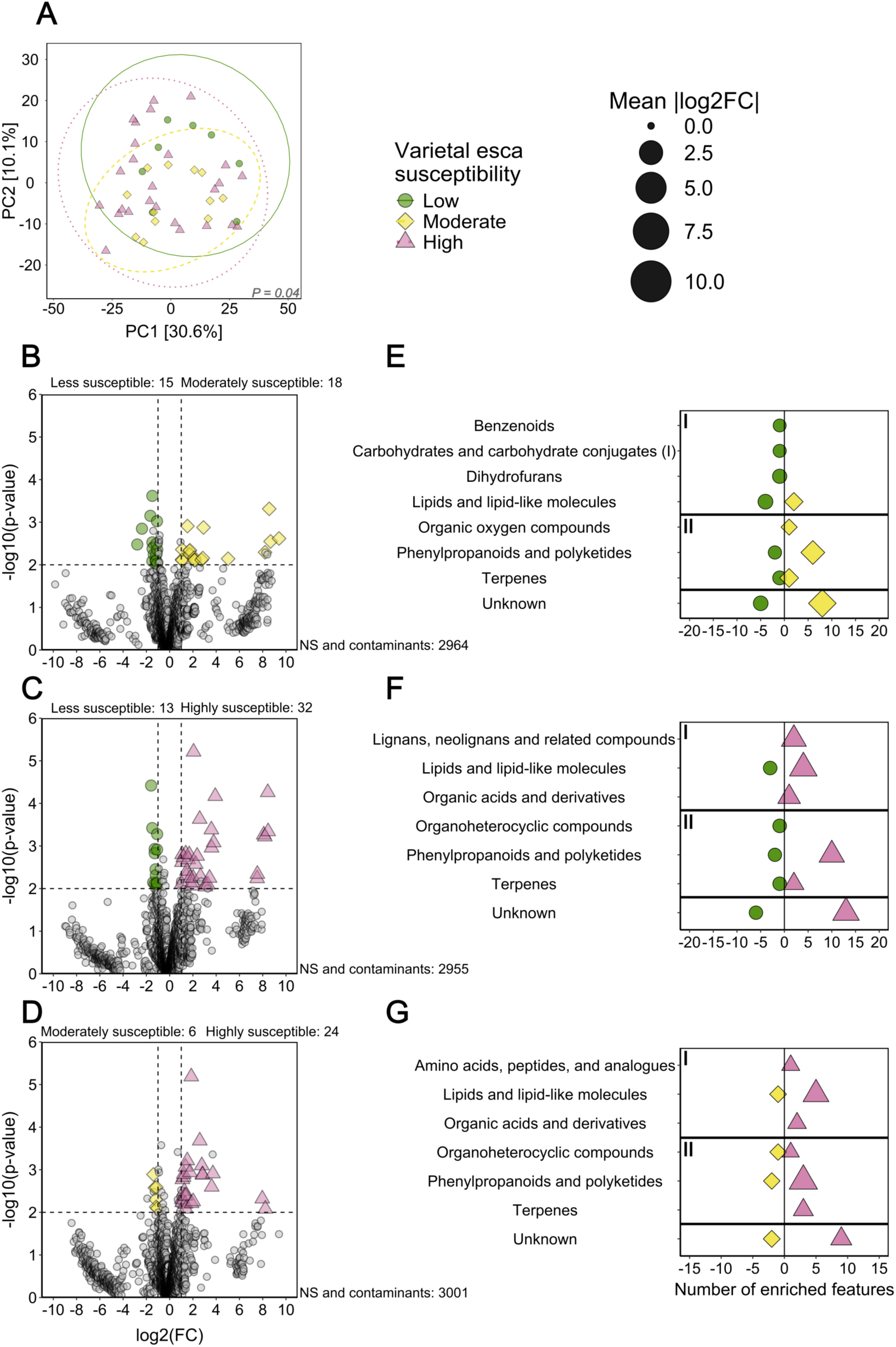
Effects of varietal susceptibility to esca on the metabolome in healthy trunk samples from plants with esca symptoms. Classes of varietal susceptibility to esca were defined on the basis of mean foliar symptom incidence and the proportion of wood displaying white-rot necrotic wood. **A,** PCA summarising the effects of varietal esca susceptibility (foliar symptoms 2017-2024, white-rot necrotic wood proportion 2024) on the structure of the trunk metabolome calculated with normalised, transformed and scaled data. Ellipses correspond to the 95% confidence interval for each group. The *P*-value for the effect of varietal esca susceptibility classes in a PERMANOVA is provided. The colours and shapes of ellipses correspond to varietal esca susceptibility classes. **B, C, D,** Volcano plots on all features. Cut-off values for significance are set at |log_2_FC| > 2 and *P* < 0.01. Non-significant (NS) and putatively contaminating features are coloured in grey. **E, F, G,** Class assignment of differentially abundant features selected through Volcano analyses (cut-off values set at |log_2_FC| > 2 and *P* < 0.01). Class assignment was performed with Classyfire and published findings. Metabolic classes are separated between primary metabolism (I, above) and secondary metabolism (II, below). Features that could not be assigned to any known compound or metabolic class are classified as “Unknown”. Panels are grouped based on pairwise comparisons across varietal esca susceptibility classes: (B, E) less (*n* = 8 plants, *n* = 5 cultivars) *vs*. moderately susceptible (*n* = 12 plants, *n* = 5 cultivars), (C, F) less *vs*. highly susceptible (*n* = 26 plants, *n* = 8 cultivars), (D, G) moderately *vs*. highly susceptible. Less susceptible cultivars are represented by green circles, moderately susceptible cultivars by yellow diamonds, and highly susceptible cultivars by red triangles.

**Fig. 6.**
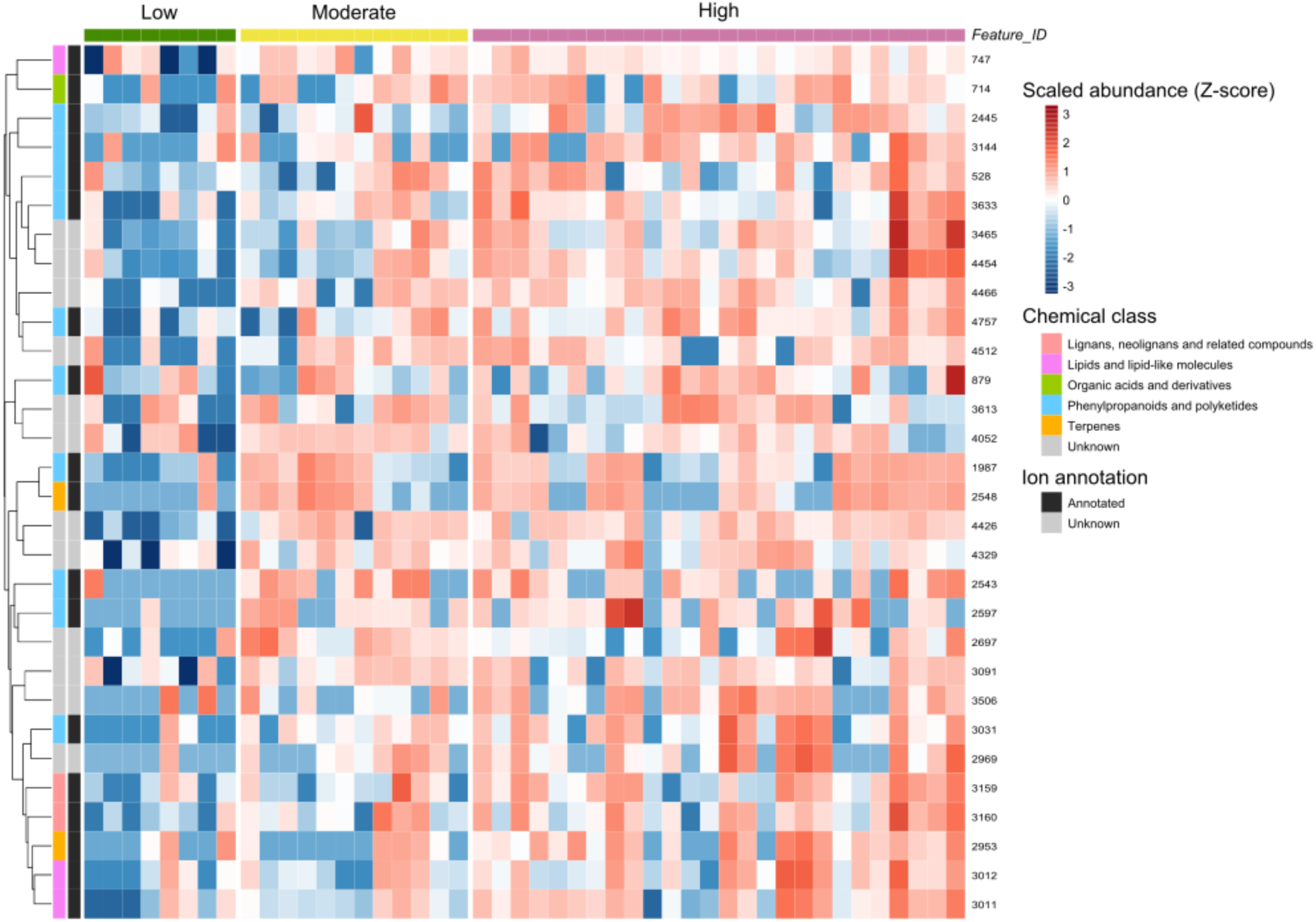
Heatmap of differentially abundant metabolites across varietal esca susceptibility classes in healthy trunk wood from symptomatic grapevines. Differentially abundant metabolites were selected from the comparison between less and highly susceptible cultivars using the same thresholds as in the volcano analyses (|log₂FC| > 2 and *P* < 0.01, without false discovery rate correction). Metabolite abundances were log₂-transformed and standardised by metabolite (Z-scores). Top 30 markers (based on |log₂FC|) were plotted and labelled according to their Feature_ID. Details on each feature are available in Supplementary Table 2. The left annotation bars indicate metabolite annotation status (annotated or unknown) and putative chemical class. Unknown features correspond to metabolites that could not be confidently assigned to a known compound. The colour scale represents standardised metabolite abundance (z-score), with blue indicating samples in which a metabolite is less abundant than its mean abundance across all samples, and red indicating samples in which it is more abundant than its mean abundance across all samples. e. Samples (columns) are grouped according to varietal esca susceptibility classes, defined from mean foliar symptom incidence (2017–2024) and the proportion of white-rot necrotic wood (2024): less susceptible (green; *n* = 8 plants, *n* = 5 cultivars), moderately susceptible (yellow; *n* = 12 plants, *n* = 5 cultivars), and highly susceptible (red; *n* = 26 plants, *n* = 8 cultivars). Metabolites (rows) are hierarchically clustered according to their abundance profiles.

Finally, to test whether constitutive metabolism also varies with cultivar susceptibility, we considered only control plants without esca symptoms. Varietal esca susceptibility slightly but significantly influenced trunk metabolic structure (Supplementary Fig. S6A). Enrichment was observed for a larger number of compounds in less susceptible than in moderately susceptible cultivars, mostly from the “phenylpropanoids and polyketides” groups (16 *vs.* 8 features; Supplementary Fig. S6B and E; *see Low vs Moderate Asymptomatic comparison* in Supplementary Table S7). Enrichment was observed for similar numbers of features in highly and less susceptible cultivars (11 *vs.* 8 features; Supplementary Fig. S6C and F; *see Low vs High Asymptomatic comparison* in Supplementary Table S7) and in moderately susceptible cultivars (9 *vs.* 12 features; Supplementary Fig. S6D and G; *see Moderate vs High Asymptomatic comparison* in Supplementary Table S7).

### Effects of cultivar and esca expression on trunk microbial communities

We obtained 3,392,501 fungal reads from 123 trunk samples, which we assigned to 365 OTUs. There was a mean number of 27,581 ± 53 fungal reads per sample (minimum: 5,035; maximum: 46,458). Five fungal phyla were retrieved: Ascomycota (87% of fungal reads), Basidiomycota (12%), a multi-affiliated fungal phylum (<1%), Oipidiomycota (<1%), and Chytridiomycota (<1%). Less than 1 % of fungal reads could not be classified at phylum level. The three most abundant fungal genera (Supplementary Fig. S7A) were *Angustimassarina* sp. (10.2% of fungal reads), *Cladosporium* sp. (9.8%) and *Alternaria* sp. (8.2%). Rarefaction curves approached saturation for most samples, indicating sufficient sampling of dominant taxa (Supplementary Fig. S8A).

We detected 709,489 bacterial reads, which were assigned to 760 OTUs. There was a mean number of 5,768 ± 31 bacterial reads per sample (minimum: 1,609; maximum: 44,977). We retrieved 17 bacterial phyla, dominated by Proteobacteria (55% of bacterial reads), Actinobacteriota (23%) and Bacteroidota (16%). The three most abundant bacterial genera (Supplementary Fig. S7B) were *Stenotrophomonas* sp. (29.6 % of bacterial reads), *Sphingomonas* sp. (6.0%) and *Methylobacterium* - *Methylorubrum* sp. (3.8%). Decontaminated OTU tables are provided in Supplementary Table S8 (ITS) and Supplementary Table S9 (16S). Rarefaction curves showed that saturation was not reached for all samples, reflecting the high diversity of bacterial communities and suggesting that rare taxa may not have been sampled (Supplementary Fig. S8B).

Consistent with the stringent filtering applied, Good’s coverage values were close to 1 across samples, reflecting a low proportion of singleton OTUs in the final dataset.

Detailed statistics for healthy trunk microbial communities are provided in Supplementary Table S10. To identify the factors influencing microbial variations, we first compared healthy wood microbial communities across varietal esca susceptibility classes (based on mean foliar symptom incidence and white-rot necrotic wood proportion) and plant esca health statuses, including cultivar as a covariate (Supplementary Tables S11 and S12). Varietal susceptibility did not significantly influence Shannon diversity index for either fungal or bacterial communities. Shannon diversity index was also not influenced by plant health status for either fungal or bacterial communities (Supplementary Fig. S9A and C). The structure of fungal and bacterial communities was significantly influenced by cultivar (23.7% explained variance, *P* = 0.001 for fungi; 21.3% explained variance, *P* = 0.001 for bacteria). Plant health status (*P* = 0.006; Supplementary Fig. S9B), varietal susceptibility (*P* = 0.001) and their interaction (*P* = 0.03) significantly influenced the structure of fungal communities but accounted for only a small proportion of the variance (respectively 1.1%, 2.4% and 1.8%). The structure of fungal communities differed slightly between highly and moderately susceptible cultivars. The structure of bacterial communities was also significantly influenced by varietal susceptibility (*P* = 0.003) but accounting for only 2.3% of the variance. Plant health status and its interaction with varietal susceptibility had no significant effect on bacterial communities (Supplementary Fig. S9D).

To understand how esca susceptibility level modulates induced modifications in wood microbial communities, we focused on plants with esca symptoms, in which varietal esca susceptibility had no significant effect on Shannon diversity index for either fungal or bacterial communities (Fig. 7A and C). Cultivar had no significant effect on Shannon diversity index for either fungal or bacterial communities. The structure of fungal and bacterial communities was significantly influenced by cultivar (36.8% explained variance, *P* = 0.002 for fungi; 36.2% explained variance, *P* = 0.007 for bacteria). However, the structure of fungal (6.3% explained variance, *P* = 0.001) and bacterial communities (5.3% explained variance, *P* = 0.04) was only slightly influenced by varietal susceptibility class. The structure of fungal communities in less susceptible cultivars differed slightly from that in moderately and highly susceptible cultivars (Fig. 7B). The structure of bacterial communities differed slightly between highly and moderately susceptible cultivars (Fig. 7D). Varietal susceptibility significantly affected the proportion of putative fungal antagonists of wood pathogens, but not of putative fungal wood pathogens (Fig. 7E) and putative bacterial antagonists of wood pathogens (Fig. 7G). Fungal antagonists, including *Cladosporium* spp., *Aureobasidium pullulans* and *Epicoccum nigrum*, were less abundant in highly susceptible cultivars with symptoms whereas the pathogen *P. chlamydospora* was more abundant in highly susceptible cultivars (Fig. 7F).

**Fig. 7.**
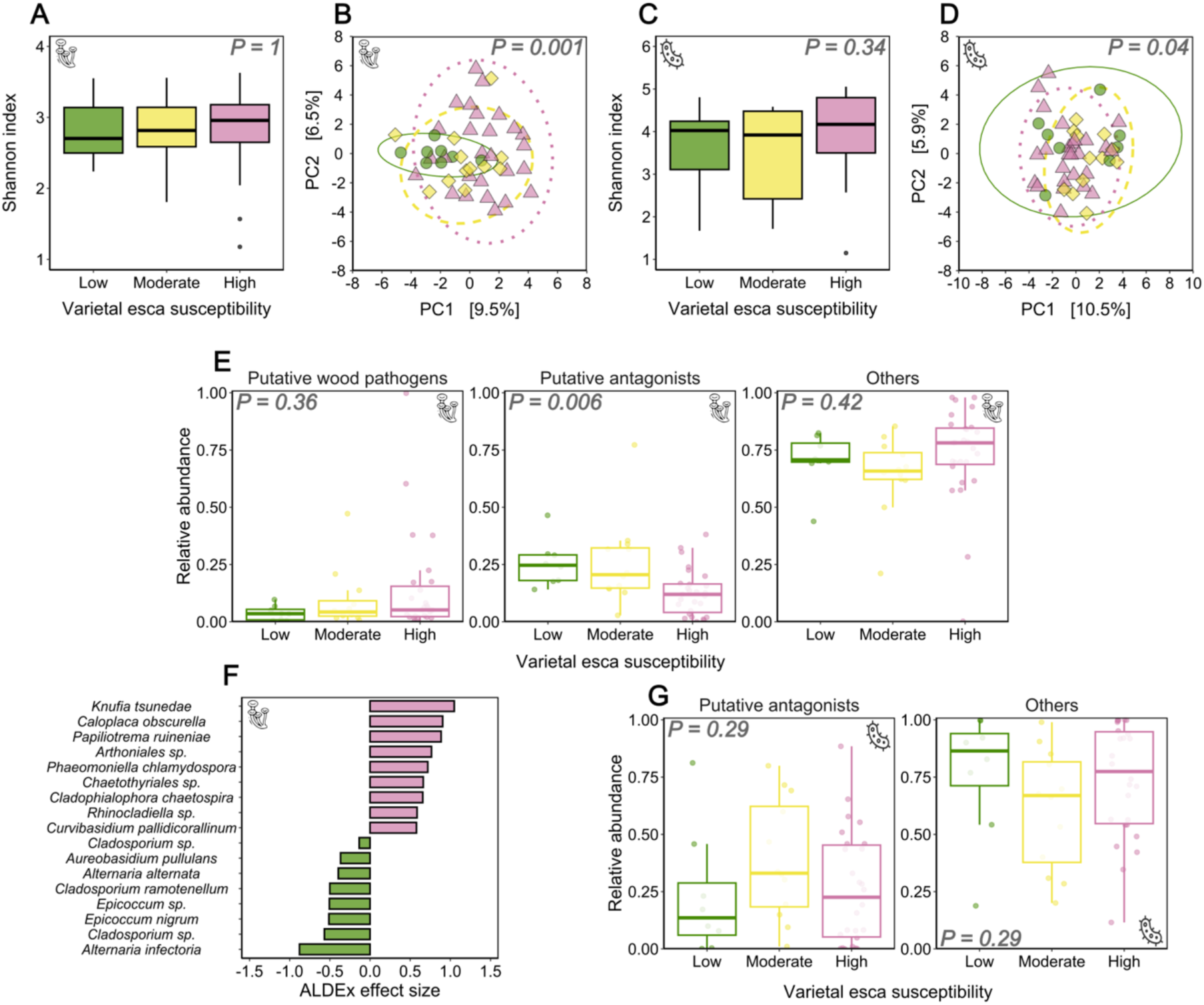
Effects of varietal esca susceptibility (*n* = 23 cultivars) on microbial communities in healthy trunk samples from plants with esca symptoms. Less susceptible cultivars (*n* = 8 plants, *n* = 5 cultivars) are shown as green circles/bars/boxplots, moderately susceptible cultivars (*n* = 12 plants, *n* = 5 cultivars) are shown as yellow diamonds/bars/boxplots, and highly susceptible cultivars (*n* = 26 plants, *n* = 8 cultivars) are shown as red triangles/bars/boxplots. We studied fungal (A, B, E, F) and bacterial (C, D, G) communities, illustrated by pictograms. **A, C,** Effect of varietal esca susceptibility (foliar symptoms 2017-2024, white-rot necrotic wood proportion 2024) on Shannon index. Boxplots display the median and interquartile range, with whiskers extending to the minimum and maximum values, excluding outliers, which are shown as individual black points. *P*-values correspond to the effects of varietal esca susceptibility in LMM analyses. **B, D,** PCA summarising the effects of varietal esca susceptibility on the structure of microbial communities calculated with CLR-transformed data. Ellipses correspond to the 95% confidence interval for each group. *P*-values correspond to the effects of varietal esca susceptibility in PERMANOVA analyses. **E, G,** Relative abundance of putative wood pathogens, putative antagonists of wood pathogens and other taxa across varietal susceptibility classes. *P*-values correspond to the effects of varietal esca susceptibility in ANOVA analyses. **F,** OTUs displaying significant differential abundance between varietal susceptibility classes according to an ALDEx2 analysis on RLE-transformed data. Features with a *p*-value < 0.05 are shown, and each bar is coloured according to the group in which the OTU is overabundant.

Finally, to test whether constitutive microbiome varies with cultivar susceptibility, we focused on control plants with no esca symptoms. Varietal esca susceptibility did not significantly influence Shannon diversity index for either fungal or bacterial communities (Supplementary Fig. S10A and C). Cultivar did not significantly influence Shannon diversity index for either fungal or bacterial communities. The structure of fungal and bacterial communities was significantly influenced by cultivar (34.5% explained variance, *P* = 0.001 for fungi; 32.3% explained variance, *P* = 0.001 for bacteria; Supplementary Fig. S10B and D). The structure of fungal communities was slightly influenced by the varietal susceptibility class (3.3 % of explained variance, *P* = 0.003), but not the structure of bacterial communities (*P* = 0.60). Varietal susceptibility had no effect on the proportions of putative fungal wood pathogens, putative fungal antagonists of wood pathogens (Supplementary Fig. S10E) and putative bacterial antagonists of wood pathogens Supplementary Fig. S10F). Complete lists of differentially abundant fungal and bacterial species, in control plants or plants with esca symptoms, are provided in Supplementary Table S13.

## Discussion

We investigated the role of wood-microorganism interactions to identify drivers of internal wood degradation and varietal susceptibility to esca across grapevine cultivars. The expression of foliar symptoms of esca in individual plants is associated with a higher proportion of internal necrosis relative to healthy wood (Maher et al., 2012; Ouadi et al., 2019; Gastou et al., 2026). We show here, across a large range of cultivars, that white-rot was the type of necrosis best correlated with foliar symptom history. However, we did not identify thresholds of white-rot necrotic wood or healthy wood proportions clearly separating control and plants with a history of esca foliar symptoms, as previously suggested (Maher et al., 2012; Gastou et al., 2026). In addition, we found that the proportion of white-rot necrotic wood in plants with a history of esca foliar symptoms was higher in susceptible cultivars suggesting a link between white-rot wood degradation and constitutive susceptibility to esca. This is a key information for predicting esca susceptibility across cultivars.

We hypothesised that cultivar differences in susceptibility to white-rot development originate from differences in the properties of healthy wood, including both plant and microbial traits. We tested three non-exclusive mechanisms that could explain this intraspecific variability: (i) susceptible cultivars differ in constitutive wood composition, resulting in greater degradability of healthy wood (Rolshausen et al., 2008; Schilling et al., 2021), (ii) susceptible cultivars differ regarding their metabolic response induced by esca, and (iii) susceptible cultivars harbour distinct healthy-wood microbial communities that influence the establishment or activity of Basidiomycota white-rot pathogens (Bekris et al., 2021).

The intraspecific variability of wood composition was low but of significant ecological and physiological relevance across 16 *V. vinifera* cultivars (same plants on which wood integrity was studied) of various origins and genetic groups. Indeed, cultivars susceptible to esca contained smaller amounts of extractives and had a slightly higher hemicellulose content (for plants with a history of esca foliar symptoms). Hemicellulose, an amorphous heterogeneous short-chain polysaccharide, is a wood component prone to degradation (Xiao et al., 2013; Pacetti et al., 2022; Haidar et al., 2024): however, we were unable to establish a strong direct link between hemicellulose content and the proportion of white-rot at the plant level. Previous studies also suggested that the amount of lignin, the most recalcitrant wood polymer, often associated with phenolic compounds, was negatively correlated with fungal degradation, even by wood pathogens, in grapevine and olive trees (Schwarze, 2007; Rolshausen et al., 2008; Markakis et al., 2019). Here, we found no significant varietal association between high total lignin content and a low proportion of white-rot necrotic wood, building on previous findings suggesting that grapevine is more susceptible to *F. mediterranea*-induced decay than beech (*Fagus sylvatica*; Schilling et al., 2022), which has a higher lignin content. As guanyl acyl-rich cell walls are generally more resistant to white-rot decay, estimating the distribution of lignin subunits (S:G ratio) in grapevine - which has not been accomplished in this study - is the next step to unravelling the role of lignin in grapevine wood decay. This can be accomplished using histological methods (e.g. Mäule or phloroglucinol staining, Pradhan Mitra and Loqué, 2014; Yamashita et al., 2016), spectroscopy (e.g., Nuclear Magnetic Resonance or Fourier-transform infrared spectroscopy; Lu et al., 2017) or wet chemistry (e.g., thioacidolysis, nitrobenzene oxidation, or hydrogenolysis; Eswaran et al., 2022).

The positive association, at the cultivar level, between the overall amount of wood extractives (*i.e.* including all secondary metabolites) and resistance to wood degradation is consistent with the properties of those compounds: antifungal or antimicrobial activities, cell wall reinforcement (Imamura, 1989; Valette et al., 2017; Schilling et al., 2021). We used untargeted metabolomics to characterise candidate low-molecular weight compounds contributing to weak cultivar susceptibility, even though it was performed on different samples - different plants, season (summer, during esca foliar symptom expression) and year. Metabolites classified as stilbenes, tannins, anthocyanins and flavonoids were among the phenylpropanoids particularly abundant in the control plants of less susceptible cultivars potentially involved in varietal tolerance: for example, stilbenoids are associated with trunk disease tolerance in grapevine (Lemaitre-Guillier et al., 2020; Khattab et al., 2021). However, caution is required in interpretation, due to the small number of compounds considered and the considerable variability between cultivars. Furthermore, the lack of comprehensive data on the same individual plants limits the scope of these trends. Thus, the protective role of some extractive compounds will now have to be experimentally confirmed. More broadly, integrating data from pathology, biochemistry, metabolomics, and microbial communities across the same set of plants in the same year represents a major avenue of research for deciphering the relationship between wood-decaying fungi, plant defense, and white-rot development.

In contrast to the significant effect of wood properties, we show not only that changes to healthy wood fungal and bacterial communities are not associated with the presence of foliar symptoms of esca - in line with previous findings (Hofstetter et al., 2012; Del Frari et al., 2019; Gastou et al., 2025) - but also that differences in microbial communities contribute little to varietal susceptibility to esca. Importantly, metabarcoding studies were not conducted on the same plants as necrosis quantification. We therefore cannot establish a causal direct relationship between white-rot proportion and the properties of microbial communities at the individual plant level, but this does not preclude noting the absence of a significant association between white-rot development and the microbiome at the cultivar level. In any case, this weak contribution (*i.e.* very low explained variance) of endophytic communities to differences between cultivars echoes the fact that they do not explain differences in esca susceptibility between vineyards planted with the same plant material (Monod et al., 2025). We found no evidence for decreases in microbial diversity or enhancement of the pathogenic component (fungal wood pathogens or wood degradation-promoting bacteria, *e.g. Paenibacillus* spp., Haidar et al., 2024), two frequent responses of the microbiome to complex diseases (Steinrucken et al., 2016; Solís-García et al., 2021). We found that cultivar is a major factor structuring grapevine trunk endophytic communities, as for other microbial compartments: rhizosphere and root endosphere (Aguilar et al., 2020; Lailheugue et al., 2024), endosphere and episphere of current-year stems (Dissanayake et al., 2018; Gastou et al., 2025), bark epiphytes (Awad et al., 2020), phyllosphere (Singh et al., 2018) or carposphere (Zhang et al., 2019). However, these constitutive varietal differences (*i.e.* in asymptomatic plants) did not determine differences in esca susceptibility between cultivars. Microbial diversity and structure, as well as wood pathogen abundance, were similar in different susceptibility classes. *F. mediterranea* relative abundance in healthy wood was lower than in previous reports (Del Frari et al., 2019; Chambard et al., 2026) and was not higher in esca-symptomatic plants. This may be due to experimental biases — we sampled functional sapwood from the current year rather than specifically taking wood at the edge of necrotic areas (as in Nerva et al., 2022). Alternatively, *F. mediterranea* may become active later in the season, or may be more abundant and active in older plants (*e.g.* 30-year-old in Chambard et al., 2026 *vs.* 15 here). The abundance of *F. mediterranea* at the time of sampling, when white-rot is already present, may not be a reliable indicator, suggesting that it should be studied earlier, before wood decay begins. Overall, this currently limits our understanding of the wood-pathogen interactions underlying the correlation between high white-rot proportion and the presence of esca foliar symptoms. Moreover, we found no relationship between other types of wood necrosis caused by wood pathogens, and esca incidence or varietal susceptibility. Black necrotic wood, classically associated with Ascomycota wood pathogens (*e.g. Phaeomoniella chlamydospora*, *Phaeoacremonium minimum*, *Botryosphaeriaceae* spp.), was similarly abundant in control and plants with a history of esca foliar symptoms, and varied between cultivars regardless of their susceptibility to esca. Interestingly, esca symptoms cannot be reproduced by artificial inoculations of Ascomycota wood pathogens (Mugnai et al., 1999) while promising results have been obtained using Basiodiomycota (Brown et al. 2019). The abundance and activity of Ascomycota pathogens does not differ significantly between asymptomatic and symptomatic vines (Del Frari et al., 2019; Chambard et al., 2026; Gastou et al., 2026). Esca foliar symptoms are inhibited under drought conditions, despite an increase in *P. chlamydospora* abundance and activity (Leal et al., 2024; Chambard et al., 2026; Gastou et al., 2026). Finally, different varietal susceptibility gradients are obtained depending on whether one considers pathogen-induced necrosis or the incidence of esca foliar symptoms (Gastou et al., 2024). These findings converge in support of the hypothesis that Ascomycete pathogens, and *P. chlamydospora* in particular, play only a limited role in esca symptom expression. *F. mediterranea*, which has the ability to degrade healthy wood directly (Moretti et al., 2023; Puca et al., 2025), would not need pioneer pathogens to cause white-rot. Conversely, the abundance of putative fungal antagonists of grapevine wood pathogens — especially *Aureobasidium pullulans* (already described as a marker of unstressed vines, Gastou et al. 2026), *Epicoccum* spp., *Cladosporium* spp. — was lower in symptomatic plants of susceptible cultivars than of less susceptible cultivars. More research is needed to clarify how these putative antagonists could limit esca expression and white-rot development in less susceptible cultivars. These findings challenge the prevailing paradigm that places the wood pathobiome - particularly its ascomycete component - at the heart of esca disease expression. This suggests that we should reconsider the role of other microorganisms, and above all, host-intrinsic factors.

Based on those findings, we propose two alternate putative scenarios underlying the association between internal and external esca symptoms: (i) wood degradation is directly responsible for leaf symptoms or (ii) white-rot accumulates as a consequence of esca-associated physiological modifications. In the first scenario, both phytotoxic metabolites (of plant and/or fungal origin) and by-products of wood degradation could be considered as possible triggers of foliar symptom expression at the start of summer, when fungal activity and sap flows are high (Claverie et al., 2020; Gastou et al., 2025). We show here that healthy trunk wood is enriched in certain secondary metabolites — phenylpropanoids (flavonoids, isoflavonoids, depsides, and depsidones) and triterpenoids — in response to esca foliar symptoms. These metabolic classes are known for their antimicrobial and antioxidant activities (Shah and Smith, 2020; Toffolatti et al., 2021). For instance, antipathogen action has been demonstrated for erythrin in lichens (*Roccella montagnei*; Selvam et al., 2022) and the isoflavone malonylglycitin in soybean (*Glycine max*; Watanabe et al., 2019). The role of these metabolites in plant defences suggests that they are of plant origin. Some of the classes of compounds displaying enrichment in the trunks of plants with esca symptoms are the same as those identified in stems and leaves (Magnin-Robert et al., 2017; Moret et al., 2021; Dell’Acqua, 2024; Gastou et al., 2025), supporting the idea of transport along the sap flow. In particular, we identified several compounds that consistently accumulated in the trunk, stems of the year and leaves, including erythrin, a phenyl benzoate, and three triterpenoids (Dell’Acqua, 2024; Gastou et al., 2025). However, as a broader metabolic response is observed in aerial organs, a specific response may also be at work in these organs, leading to the simultaneous or delayed accumulation of defence compounds *in situ* in aerial organs. We also identified several glycosylated phenylpropanoids that specifically accumulated in the trunk in response to esca foliar symptoms and were more abundant in susceptible cultivars. The accumulation of glycosylated phenylpropanoids was also observed in stems displaying esca symptoms in highly susceptible varieties (Gastou et al., 2025). It may reflect an inability of susceptible genotypes to produce bioactive non-glycosylated phenylpropanoids efficiently, or an attempt to detoxify overabundant phytotoxic compounds potentially occurring in such genotypes (Le Roy et al., 2016; Gastou et al., 2025). In this scenario, these non-bioactive molecules could act as a signal triggering the onset of esca symptoms, unlike certain protective bioactive compounds — mentioned above — acting as an antimicrobial barrier.

In the second scenario, we propose that white-rot necrotic wood results from physiological changes induced by esca expression. Despite its overall association with plant esca history, white-rot necrotic wood was absent from almost a third of the symptomatic plants studied. Interestingly, all the plants with no visible white-rot necrotic wood expressed esca symptoms in only one year and half of those with esca symptoms in only one year had no white-rot. This result may suggest that the presence white-rot necrotic wood is not required for esca leaf symptom expression, unlike vascular occlusions and brown stripe, which are always found in symptomatic plants (Lecomte et al., 2012; Bortolami et al., 2019, 2021a; Dell’Acqua et al., 2024). In this case, a putative increase in *F. mediterranea* abundance and activity (as shown by Chambard et al. (2026) in healthy wood) in response to esca would increase wood degradation, especially in susceptible cultivars. This increase in the amount of white-rot necrotic wood and the potentially associated pathogen load may have a retroactive effect, promoting leaf symptom expression in subsequent years. Our understanding of the mechanistic relationship between necrosis and aerial symptoms might be improved by systematically (from the graft union to the top of the arms) phenotyping wood integrity and *F. mediterranea* activity as soon as leaf symptoms appear on previously asymptomatic vines, to determine whether white-rot necrotic wood was present before symptom onset. Moreover, characterizing the virulence of *F. mediterranea* isolates originating from different cultivars — which has not been done here — would be a promising avenue of research, using approaches such as inoculation on sawdust (Schilling et al., 2022) or genomics (e.g. targeting virulence factors; Massonnet et al., 2018; Garcia et al., 2024).

## Conclusion

These results demonstrate the tight association between a single type of wood necrosis — white-rot — and varietal susceptibility to esca. By analysing both plant and microbiome traits, we uncovered marked cultivar-specific differences in wood composition. However, a low proportion of the variance was explained by microbial community structure and wood pathogen abundance. These findings support a wood-centred view of esca pathogenesis, in which plant-intrinsic properties, rather than pathogen identity, play a dominant role in disease development. Host physiology should thus be better taken into account when assessing the susceptibility of perennial plants to complex diseases, as it can modulate symptom expression under equivalent pathogen pressures. It seems now essential to characterize the dynamics of defense compound production and wood degradation over time and across different organs within the same plants. Overall, these results indicate that the choice of variety, together with pruning practices, can be used to limit necrosis developments (Lecomte et al., 2018; Kraus et al., 2022) with white-rot removal by trunk surgery as a last resort (often efficient, but invasive, expensive and sometimes unsuccessful, Pacetti et al., 2021; Lecomte et al., 2022; Dewasme et al., 2023), mitigating esca expression and increasing vineyard sustainability.

## Supplementary data

**Supplementary Table S1.** List of the 23 grapevine cultivars included in this study.

**Supplementary Table S2.** Effects of cultivar, organ, plant esca history, their interactions and the plant health status on the distribution of wood types.

**Supplementary Table S3.** Effects of cultivar, organ, the number of symptomatic years and their interactions on the distribution of wood types in plants with a history of esca foliar symptoms.

**Supplementary Table S4.** Effects of cultivar, organ, plant health status and their interactions on the distribution of wood types in plants with a history of esca foliar symptoms.

**Supplementary Table S5.** Effects of cultivar, plant esca history, their interaction and plant health status on the distribution of wood components.

**Supplementary Table S6.** Effects of proportions of each wood component, cultivar, plant esca history and their interaction on the proportion of white-rot necrotic wood in the upper trunk cross-section.

**Supplementary Table S7.** List of metabolic features with differential abundances across plant health statuses and varietal susceptibility clusters in healthy trunk samples.

**Supplementary Table S8.** OTU table corresponding to the entire decontaminated ITS metabarcoding dataset for healthy trunk samples.

**Supplementary Table S9.** OTU table corresponding to the entire decontaminated 16S metabarcoding dataset for healthy trunk samples.

**Supplementary Table S10.** Effects of plant health status, varietal susceptibility cluster and cultivar on Shannon index and beta-diversity in healthy trunk microbial communities.

**Supplementary Table S11.** Mean ± SEM of Shannon index for fungal and bacterial communities in healthy trunk samples across cultivars and plant health statuses.

**Supplementary Table S12.** Cultivar effect on the beta-diversity of fungal and bacterial communities in healthy trunk samples from plants with and without esca symptoms.

**Supplementary Table S13.** Summary of the results obtained in Aldex2 differential analyses on healthy trunk samples.

**Supplementary Figure S1.** Details of wood sampling and visual determination of the index of black punctate necrosis.

**Supplementary Figure S2.** K-means clustering and silhouette analysis of grapevine cultivars based on esca-related traits.

**Supplementary Figure S3.** Mean distribution of healthy and necrotic wood across plant esca histories and cultivars.

**Supplementary Figure S4.** Effect of plant esca history on the mean proportion of visually altered sapwood perimeter and the index of black punctate necrosis.

**Supplementary Figure S5.** Effects of esca expression on the metabolome in healthy trunk samples (*n* = 123 plants, *n* = 23 cultivars).

**Supplementary Figure S6.** Effects of varietal susceptibility to esca on the metabolome in healthy trunk samples from asymptomatic plants.

**Supplementary Figure S7**. Distribution of the 20 most abundant taxa in grapevine healthy trunk samples.

**Supplementary Fig. S8.** Rarefaction curves of microbial communities (fungi and bacteria).

**Supplementary Figure S9.** Effects of plant health status on microbial communities in healthy trunk samples (*n* = 123 plants, *n* = 23 cultivars).

**Supplementary Figure S10.** Effects of varietal esca susceptibility (*n* = 23 cultivars) on microbial communities in healthy trunk samples from asymptomatic plants.

## Acknowledgements

We thank Agnes Destrac Irvine and Cornelis van Leeuwen (EGFV, Univ. Bordeaux, Bordeaux Sciences Agro, INRAE, ISVV) for providing access to the VitAdapt experimental vineyard. We also thank the Experimental Viticultural Unit of Bordeaux 1442, INRAE, F-33883 Villenave d’Ornon for vineyard management. We thank Sylvie Bastien (SAVE), Orlane Touzet (UMR BioForA, Phenobois) and Audrey Morin (SAVE) for their contributions to experiments. We thank Samuele Moretti and Corinne Vacher for useful scientific and technical advice. We thank Alexandre Chataigner (SAVE) and Linda Stammitti-Bert (EGFV) for their support with molecular biology, Manon Chargy and Paola Fournier for providing microbial strains used as positive controls. We also thank the Genome Transcriptome facility of Bordeaux (PGTB, Cestas, France).

## Author contributions

C.E.L.D. and P.G. designed the experiment; L.A., T.C., P.G., G.C., N.F. and C.E.L.D obtained trunk cross-sections and performed wood sampling; T.C. analysed images of trunk cross-sections; N.B. performed chemical analyses to determine wood composition; P.G. and N.F. sampled healthy wood in the field and ground the samples; C.R., P.P. and the MetaboHUB-Bordeaux team performed untargeted metabolomic experiments (extraction, LC-MS and MS-DIAL analysis); P.P. contributed to metabolomic data analysis and interpretation; P.G. and N.F. performed DNA extraction and PCR for metabarcoding; P.G. analysed all datasets, produced the figures and wrote the first version of the paper under the supervision of C.E.L.D.; all the authors contributed to scientific discussion, and critically revised and approved the final version of the manuscript.

## Conflict of interest

The authors have no competing interests to declare.

## Funding

Pierre Gastou’s PhD grant was awarded by the French *Ministère de l’Enseignement Supérieur et de la Recherche*. This study received financial support from Château-Figeac (Saint-Emilion), the French government in the framework of the IdEX Bordeaux University “Investments for the Future” programme / GPR Bordeaux Plant Sciences, and the French National Research Agency (ANR) in the framework of the “Investments for the Future Programme”, within the Cluster of Excellence COTE (ANR-10-LABX-45). Some experiments (ITS and 16S sequencing) were performed at the Genome 706 Transcriptome Facility of Bordeaux (Grants from *Investissements d’avenir, Convention 707 attributive d’aide EquipEx Xyloforest* ANR-10-EQPX-16-01).

## Data availability

All sequencing datasets (ITS and 16S metabarcoding) are available in the ENA database under accession number PRJEB100545. Datasets used for statistical analyses (processed metabolomics data, biochemical data, wood type quantifications) are available in the INRAe dataverse: https://doi.org/10.57745/YWR2PC, Recherche Data Gouv, V1

## Notes

https://doi.org/10.57745/YWR2PC

### Competing Interest Statement

The authors have declared no competing interest.

### Summary of Updates

Corrections have been made to the title and abstract; the description of the methods to make them more concise; and various minor changes in the body of the text. One section has been deleted (pathogen inoculation trials).

