## Supplementary figures for "Contrasting wood chemical traits are associated with differences in wood degradation and esca susceptibility among grapevine cultivars"

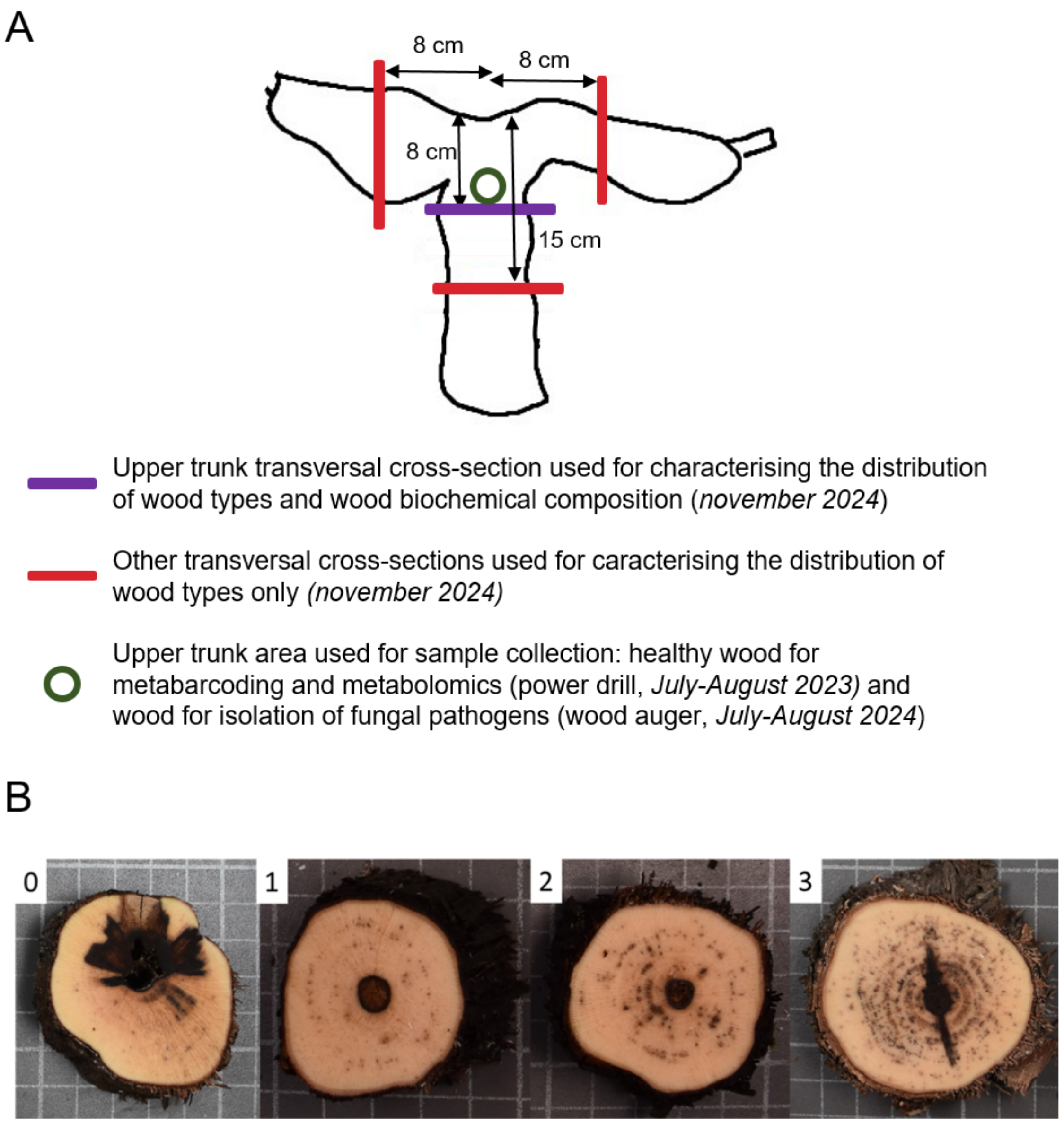


**Supplementary Fig. S1. Details of wood sampling and visual determination of the index of black punctate necrosis. A,** Conceptual representation of wood samplings performed on grapevine arms and trunks in 2023 and 2024. **B,** Qualitative scale for visually characterising the density of black punctate necrosis on transversal cross-sections. It ranges from 0 (very few punctuations) to 3 (many punctuations). The following examples are presented: a section of Semillon trunk [0], a section of Ugni Blanc trunk [1], a section of Ugni Blanc trunk [2], a section of Cot trunk [3].

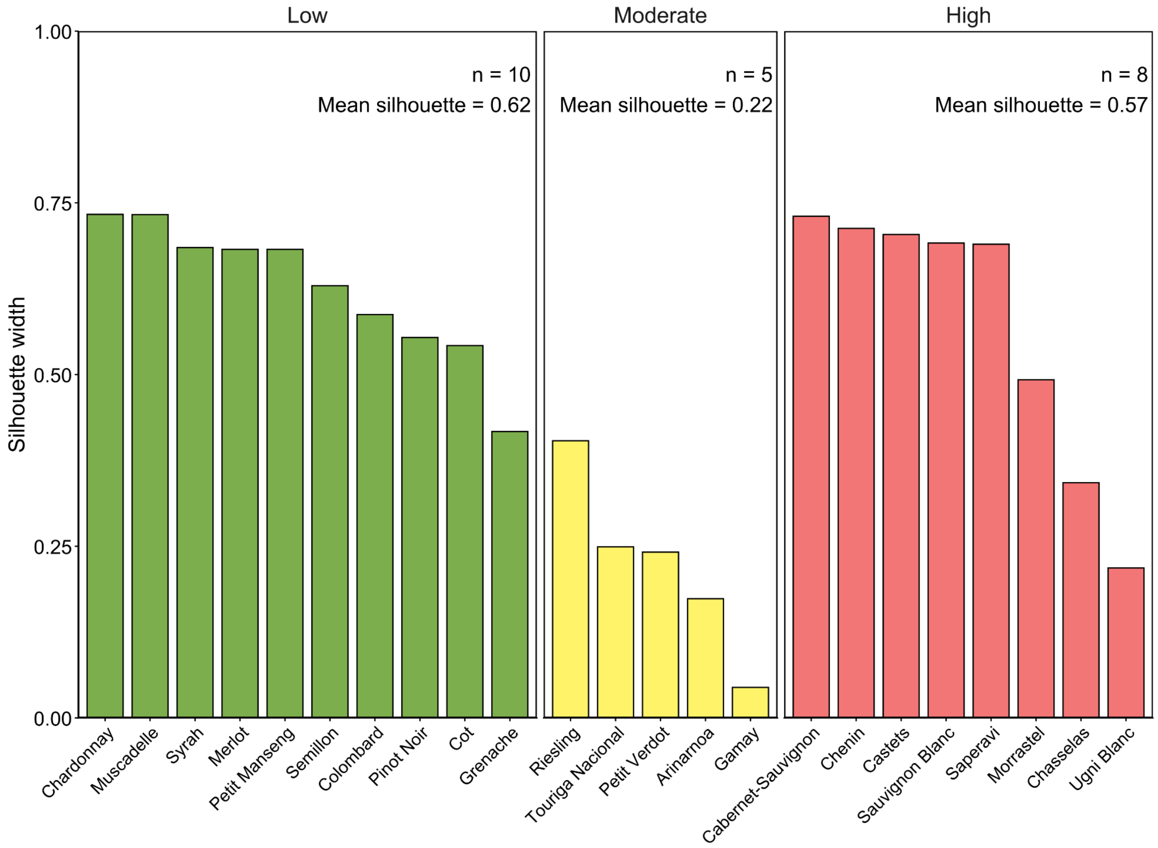


**Supplementary Fig. S2. K-means clustering and silhouette analysis of grapevine cultivars based on esca-related traits.** Cultivars were grouped using k-means clustering (k = 3) on mean foliar symptom incidence (2017-2024) and mean proportion of white-rot necrotic wood in plants with a history of esca foliar symptoms (2024). Silhouette widths quantify the quality of cluster assignment for each cultivar, with higher values indicating better within-cluster similarity. Bars are ordered by cluster and silhouette width, and cluster-level means and sample sizes are indicated.

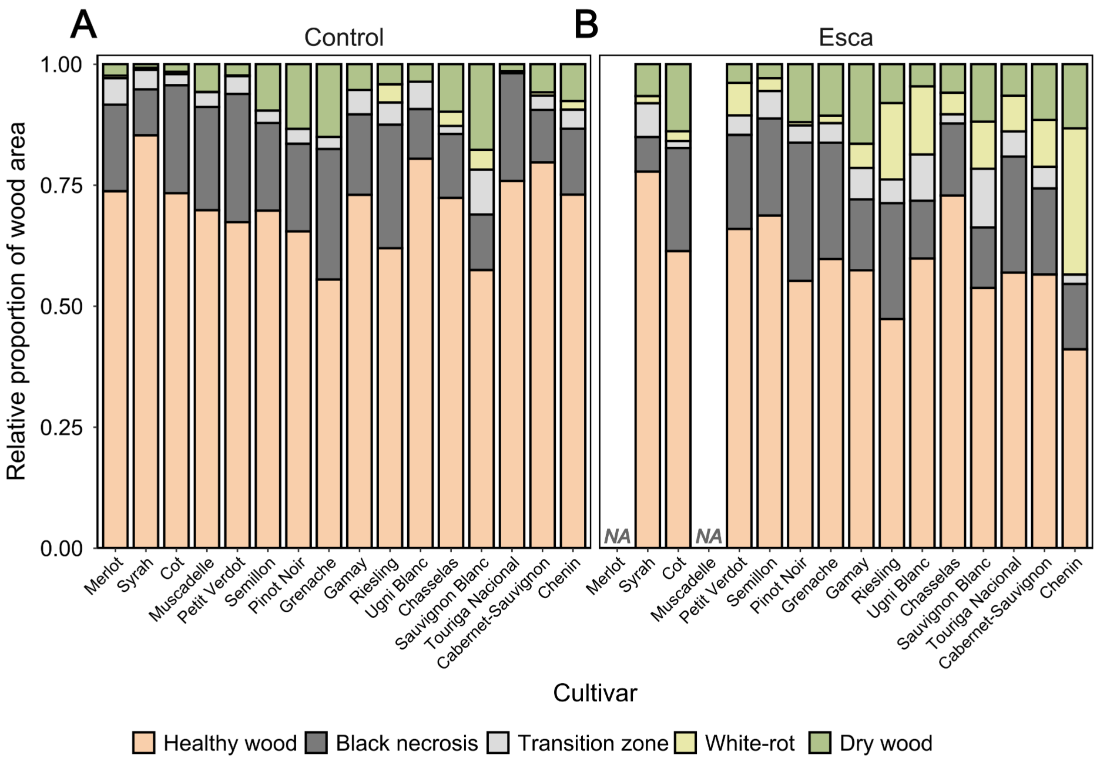


**Supplementary Fig. S3. Mean distribution of healthy and necrotic wood across plant esca histories and cultivars. A,** Control plants. **B,** Plants with a history of esca foliar symptoms. Values correspond to the mean obtained from arm (n = 191 sections) and trunk samples (n = 195 sections). The bars are coloured according to the wood type. Detailed statistics are given in Supplementary Table S3.

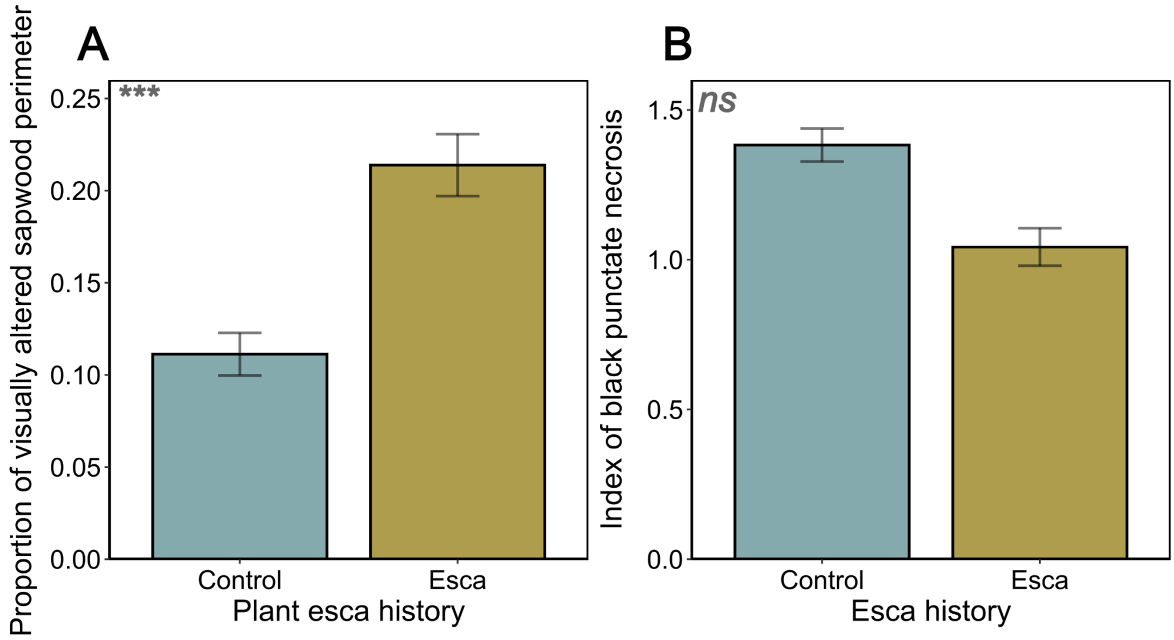


**Supplementary Fig. S4. Effect of plant esca history on the mean proportion of visually altered sapwood perimeter and the index of black punctate necrosis. A,** Proportion of visually altered sapwood perimeter (mean ± SEM). **B,** Index of black punctate necrosis (mean ± SEM) for control (n = 222 sections, n = 55 plants, n = 16 cultivars) and plants that previously displayed esca symptoms (n = 164 sections, n = 41 plants, n = 14 cultivars). Values correspond to the mean obtained from arm (n = 191 sections) and trunk samples (n = 195 sections). P-values correspond to plant esca history effects in independent Linear Mixed Models. ns: non significant, ***: P < 0.001. Detailed statistics are given in Supplementary Table S3.

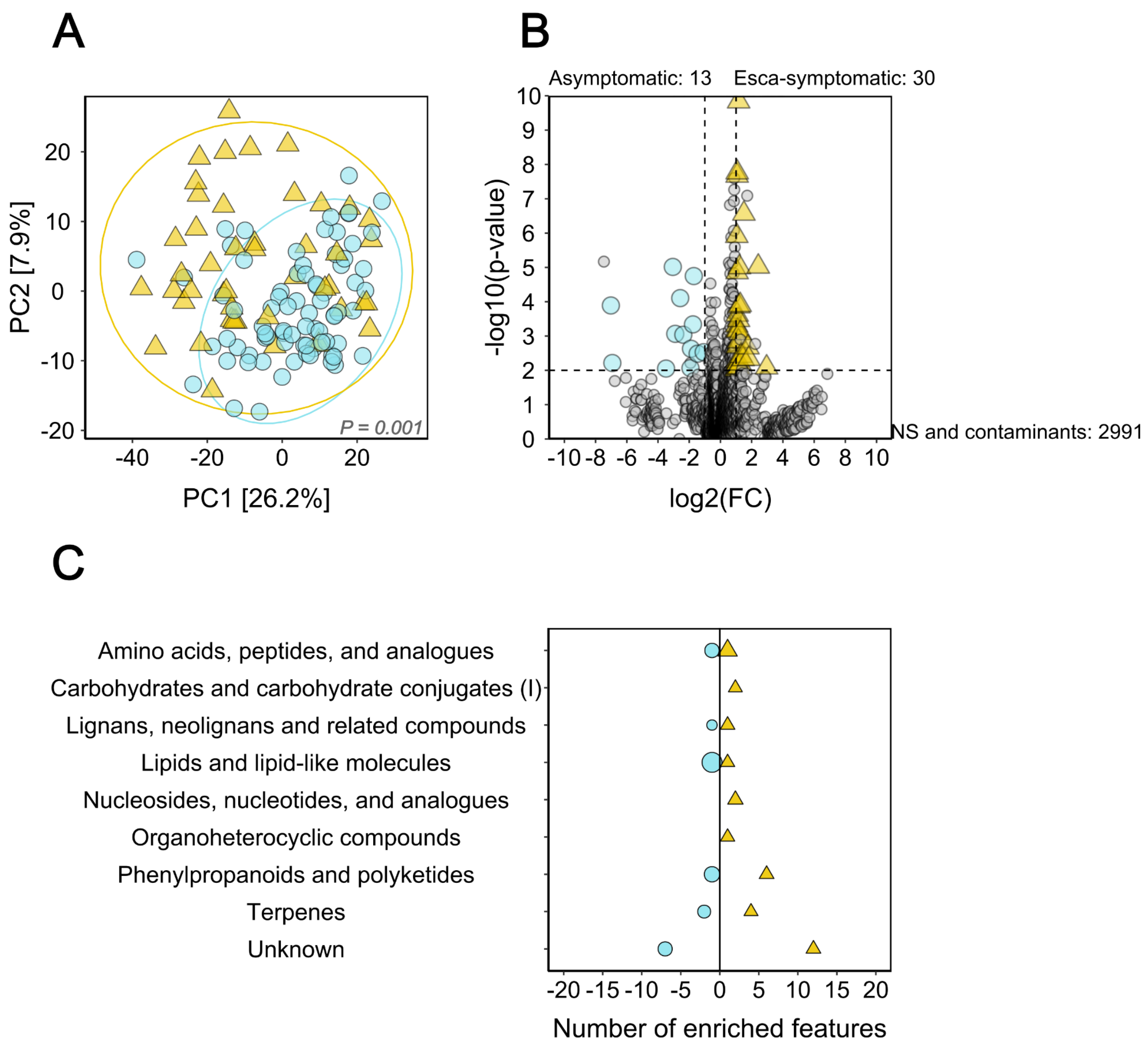


**Supplementary Fig. S5. Effects of esca expression on the metabolome in healthy trunk samples (*n* = 123 plants, *n* = 23 cultivars). A,** PCA summarising the effects of plant health status on the structure of the trunk metabolome calculated with normalised, transformed and scaled data. Ellipses correspond to the 95% confidence interval for each group. P-value corresponds to the effect of plant health status in a PERMANOVA. The colors and shapes of dots and ellipses correspond to plant health status. **B,** Volcano plots on all features. Cut-off values for significance are set at |log2FC| > 2 and P < 0.01. Non-significant (NS) and putatively contaminant features are coloured in grey. **C,** Class assignment of features displaying significant differential abundance selected through Volcano analyses (cut-off values set at |log2FC| > 2 and P < 0.01). Class assignment was performed using Classyfire and literature. Features that could not be assigned to any known compound or metabolic class are classified as “Unknown”.

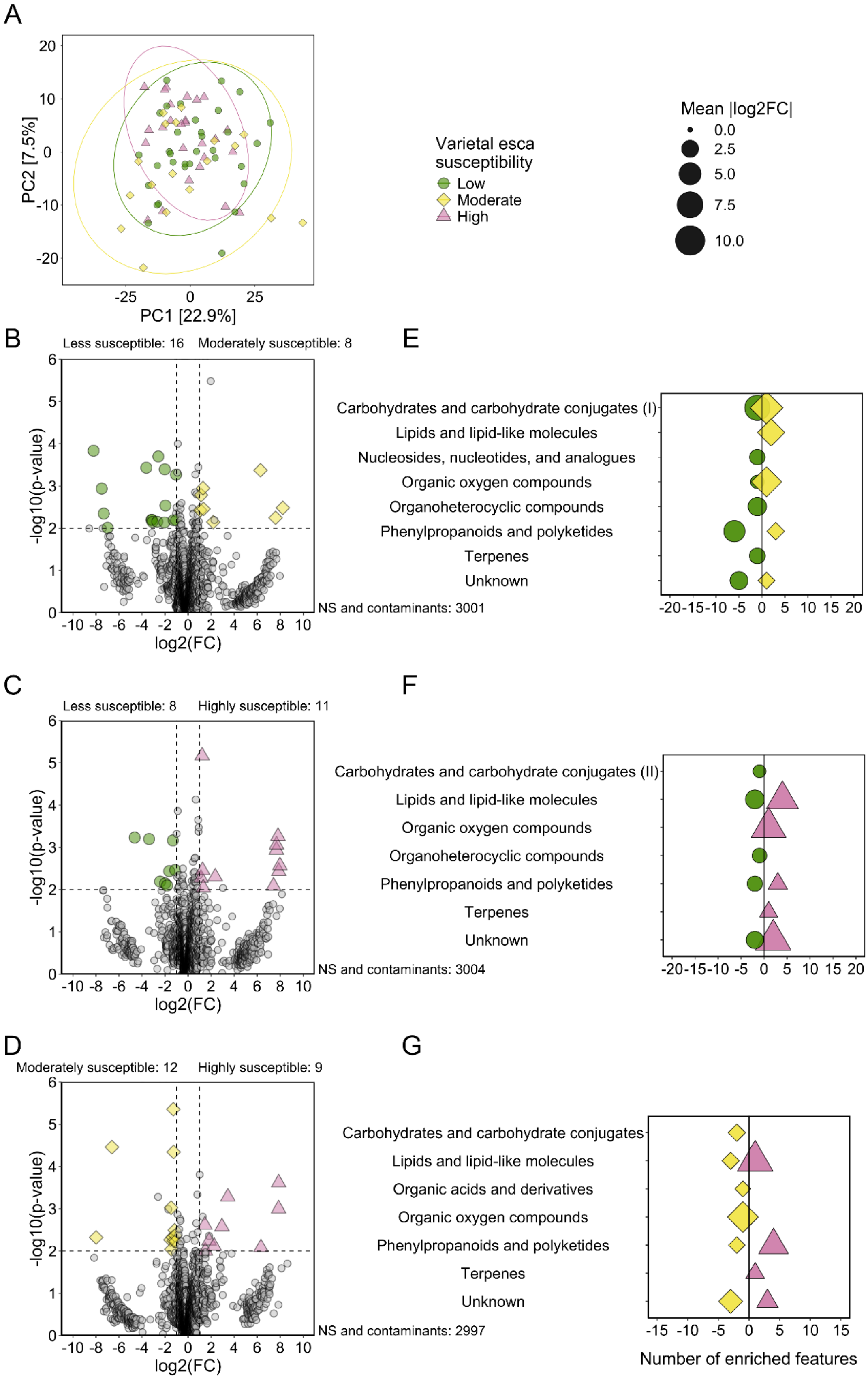


**Supplementary Fig. S6. Effects of varietal susceptibility to esca on the metabolome in healthy trunk samples from asymptomatic plants.** Classes of varietal susceptibility to esca are created based on mean foliar symptom incidence and white-rot necrotic wood proportion. **A,** PCA summarising the effects of varietal esca susceptibility (foliar symptoms 2017-2024, white-rot proportion 2024) on the structure of the trunk metabolome calculated with normalised, transformed and scaled data. Ellipses correspond to the 95% confidence interval for each group. P-value corresponds to the effect of varietal esca susceptibility classes in a PERMANOVA. The colors and shapes of dots and ellipses correspond to varietal esca susceptibility classes. **B, C, D,** Volcano plots on all features. Cut-off values for significance are set at $|\log2FC| > 2$ and $P < 0.01$. Non-significant (NS) and putatively contaminant features are coloured in grey. **E, F, G,** Class assignment of displaying significant differential abundance selected through Volcano analyses (cut-off values set at $|\log2FC| > 2$ and $P < 0.01$). Class assignment was performed using Classyfire and literature. Features that could not be assigned to any known compound or metabolic class are classified as "Unknown". Panels are grouped based on pairwise comparisons across varietal esca susceptibility classes: (B, E) less (n = 33 plants, n = 10 cultivars) *vs*. moderately susceptible (n = 18 plants, n = 5 cultivars), (C, F) less *vs*. highly susceptible (n = 26 plants, n = 8 cultivars), (D, G) moderately *vs*. highly susceptible. Less susceptible cultivars are represented by green circles, moderately susceptible cultivars by yellow diamonds, and highly susceptible cultivars by red triangles.

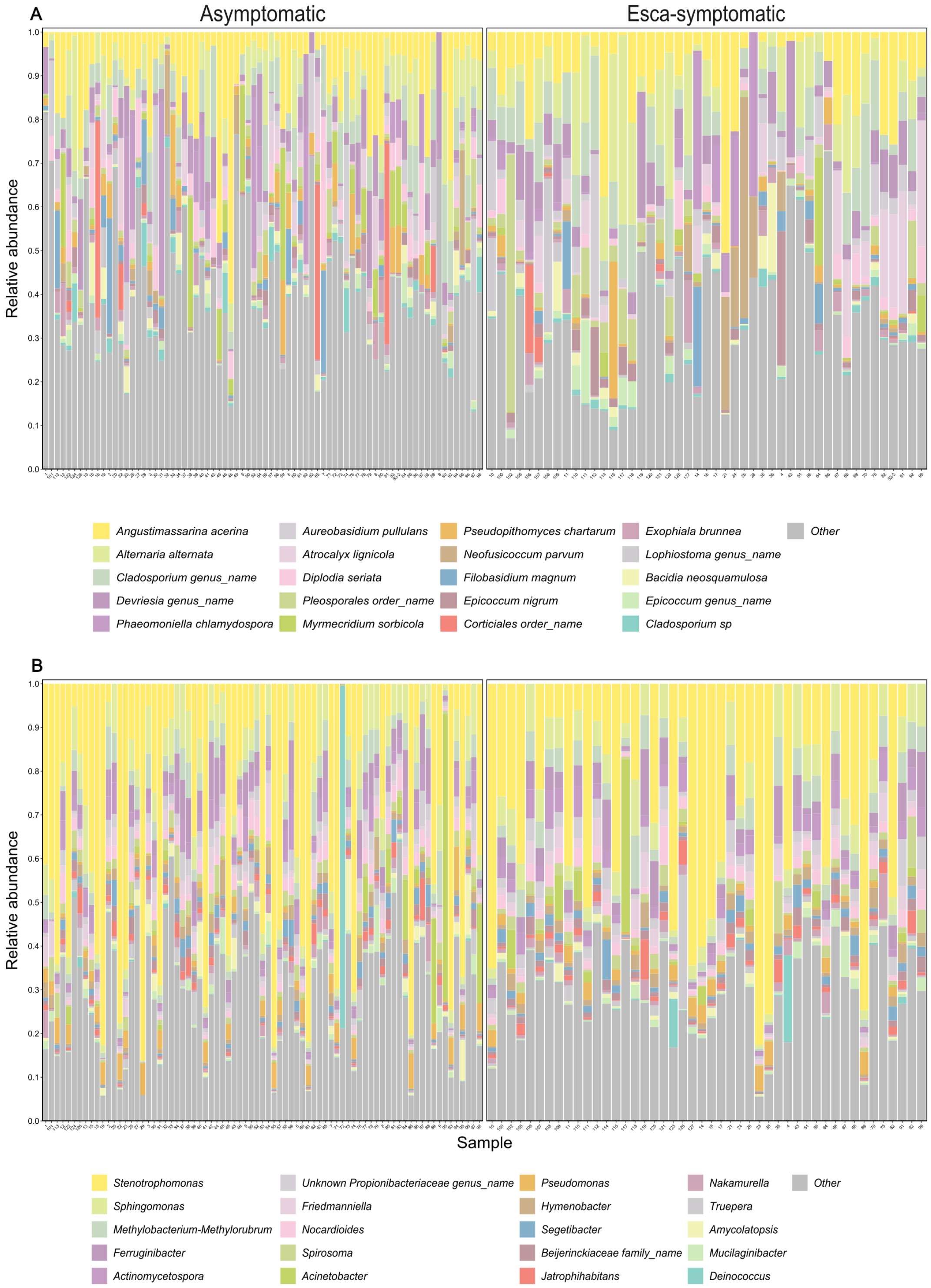


**Supplementary Fig. S7. Distribution of the 20 most abundant taxa in grapevine healthy trunk samples. A,** Distribution of fungal species (ITS metabarcoding) across plant health statuses. **B,** Distribution of bacterial genera (16S metabarcoding) across plant health statuses. Mean relative abundances are calculated for each taxa. The stacked bars are coloured according to the taxa. Less abundant taxa are grouped in the category “Other” and coloured in grey.

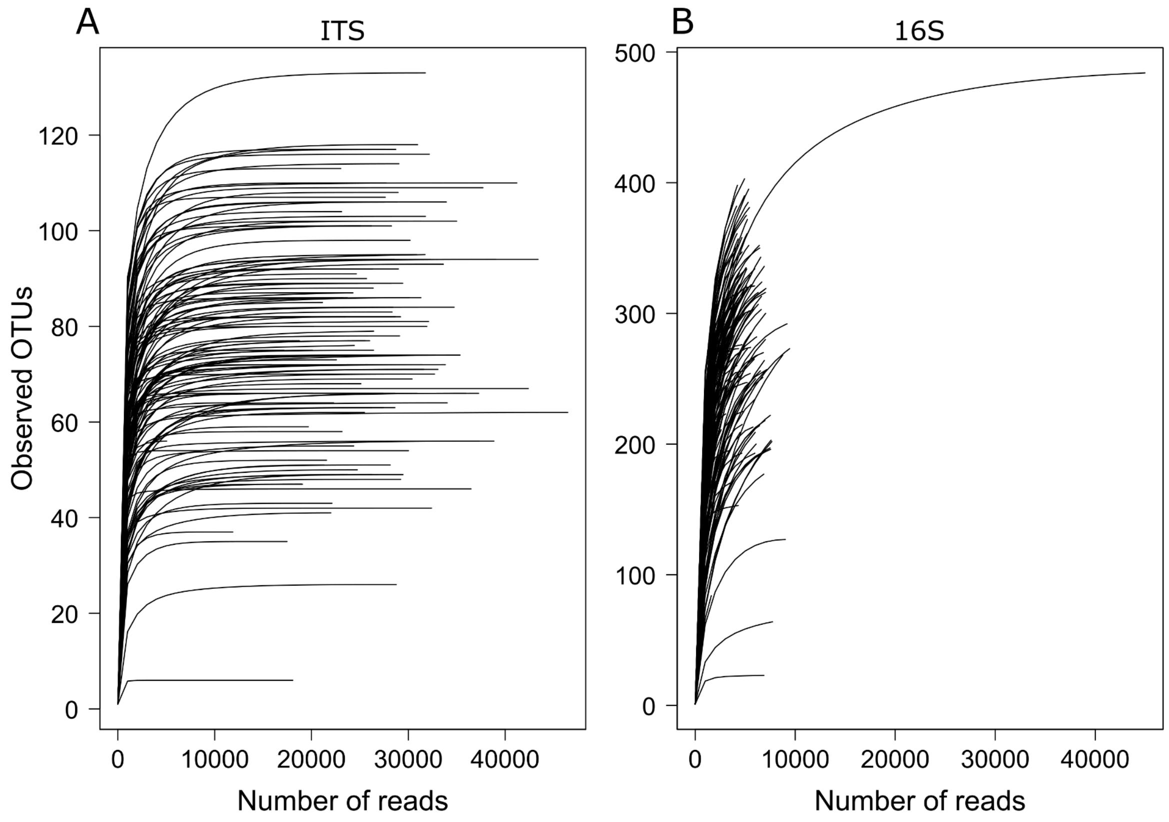


**Supplementary Fig. S8. Rarefaction curves of microbial communities (fungi and bacteria).** Rarefaction curves showing observed OTU richness as a function of sequencing depth for **(A)** ITS and **(B)** 16S datasets.

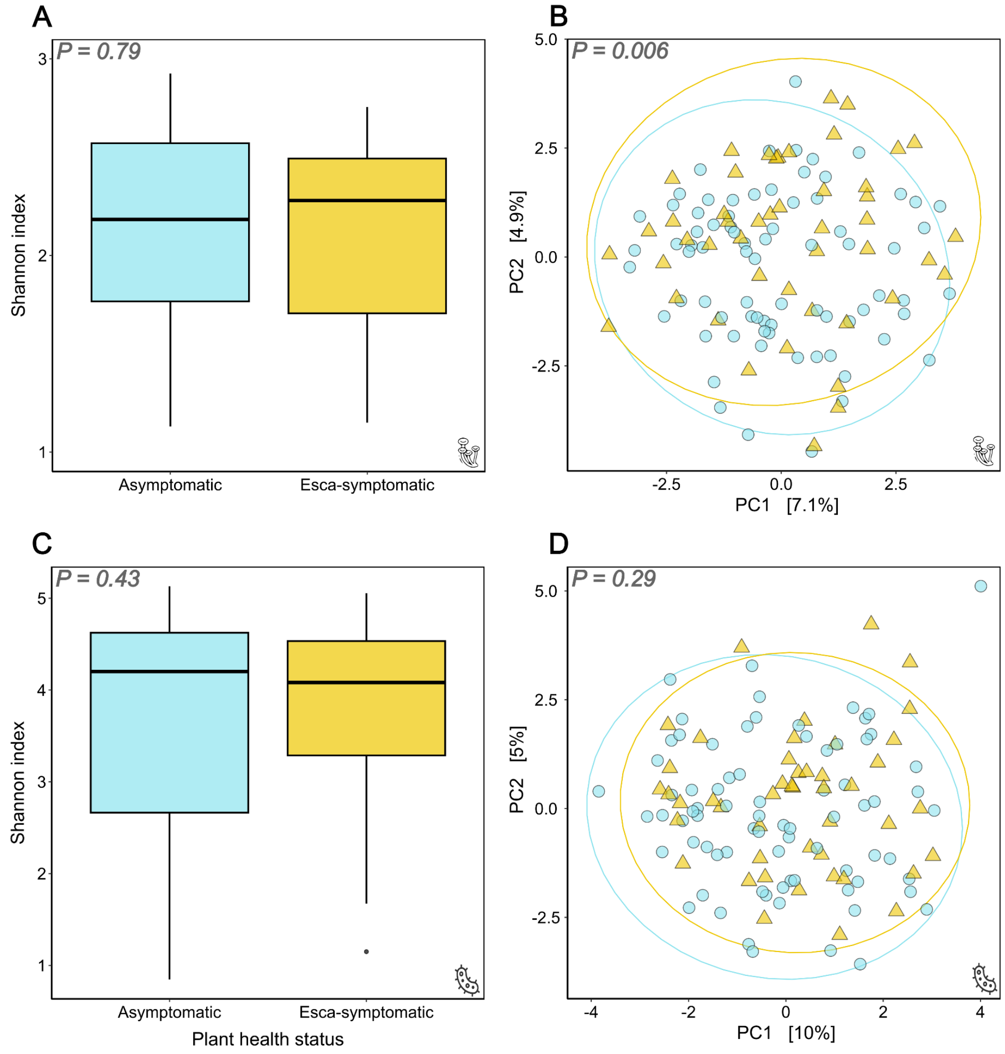


**Supplementary Fig. S9. Effects of plant health status on microbial communities in healthy trunk samples (*n* = 123 plants, *n* = 23 cultivars).** We studied fungal (A, B) and bacterial (C, D) communities as illustrated by pictograms. **A, C,** Effect of plant health status on Shannon index. Boxplots display the median and interquartile range, with whiskers extending to the minimum and maximum values, excluding outliers, which are shown as individual black points. P-values correspond to plant health status effects in LMM analyses. **B, D,** PCA summarising the effects of varietal esca susceptibility on the structure of microbial communities calculated with CLR-transformed data. Ellipses correspond to the 95% confidence interval for each group. P-values correspond to plant health status effects in PERMANOVA analyses.

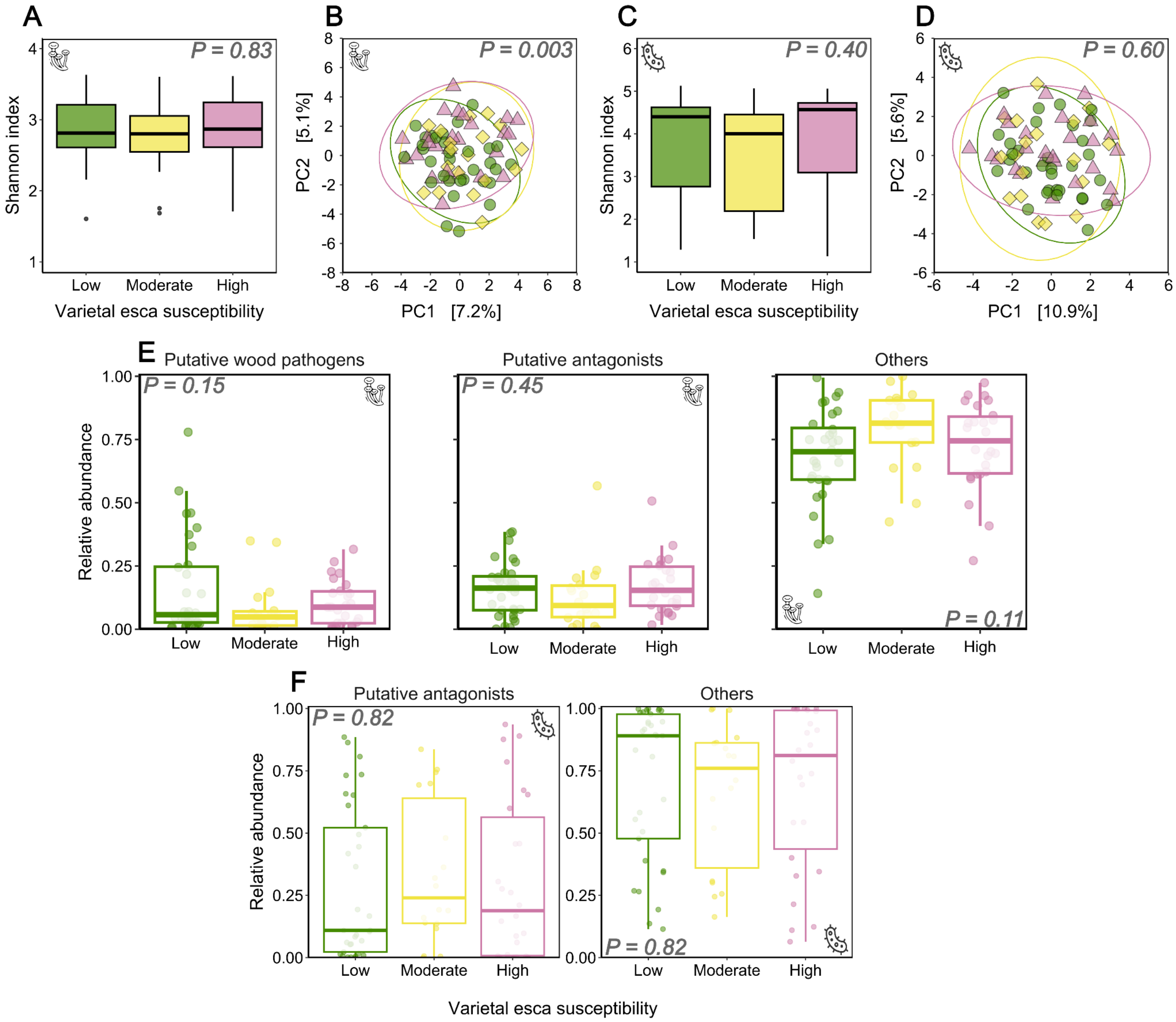


**Supplementary Fig. S10. Effects of varietal esca susceptibility (*n* = 23 cultivars) on microbial communities in healthy trunk samples from asymptomatic plants.** Less susceptible cultivars (n = 33 plants, n = 10 cultivars) are shown as green circles/bars/boxplots, moderately susceptible cultivars (n = 18 plants, n = 5 cultivars) are yellow diamonds/bars/boxplots, and highly susceptible cultivars (n = 26 plants, n = 8 cultivars) are red triangles/bars/boxplots. We studied fungal (A, B, E) and bacterial (C, D, F) communities as illustrated by pictograms. **A, C,** Effect of varietal esca susceptibility (foliar symptoms 2017-2024, white-rot proportion 2024) on Shannon index. Boxplots display the median and interquartile range, with whiskers extending to the minimum and maximum values, excluding outliers, which are shown as individual black points. P-values correspond to varietal esca susceptibility effects in LMM analyses. **B, D,** PCA summarising the effects of varietal esca susceptibility on the structure of microbial communities calculated with CLR-transformed data. Ellipses correspond to the 95% confidence interval for each group. P-values correspond to varietal esca susceptibility effects in PERMANOVA analyses. **E, F,** Relative abundance of putative wood pathogens, putative antagonists of wood pathogens and other taxa across varietal susceptibility classes. P-values correspond to varietal esca susceptibility effects in ANOVA analyses.
